# Hyperspectral Reflectance-Derived Relationship Matrices for Genomic Prediction of Grain Yield in Wheat

**DOI:** 10.1101/389825

**Authors:** Margaret R. Krause, Lorena González-Pérez, José Crossa, Paulino Pérez-Rodríguez, Osval Montesinos-López, Ravi P. Singh, Susanne Dreisigacker, Jesse Poland, Jessica Rutkoski, Mark Sorrells, Michael A. Gore, Suchismita Mondal

**Affiliations:** Plant Breeding and Genetics Section, School of Integrative Plant Science, Cornell University, Ithaca, New York, 14853, USA; Global Wheat Program, International Maize and Wheat Improvement Center (CIMMYT), Ciudad de México, 06600, México; Colegio de Postgraduados, CP 56230, Montecillos, Edo. de México; Facultad de Telemática, Universidad de Colima, Colima, Colima, 28040, México; Department of Plant Pathology, Kansas State University, Manhattan, Kansas, 66506, USA; International Rice Research Institute (IRRI), DAPO Box 7777, Metro Manila, 1301, Philippines

**Keywords:** hyperspectral reflectance, genomic prediction, high throughput phenotyping, wheat breeding, genotype-by-environment interaction, GBLUP

## Abstract

Hyperspectral reflectance phenotyping and genomic selection are two emerging technologies that have the potential to increase plant breeding efficiency by improving prediction accuracy for grain yield. Hyperspectral cameras quantify canopy reflectance across a wide range of wavelengths that are associated with numerous biophysical and biochemical processes in plants. Genomic selection models utilize genome-wide marker or pedigree information to predict the genetic values of breeding lines. In this study, we propose a multi-kernel GBLUP approach to genomic selection that uses genomic marker-, pedigree-, and hyperspectral reflectance-derived relationship matrices to model the genetic main effects and genotype × environment (*G* × *E*) interactions across environments within a bread wheat (*Triticum aestivum* L.) breeding program. We utilized an airplane equipped with a hyperspectral camera to phenotype five differentially managed treatments of the yield trials conducted by the Bread Wheat Improvement Program, International Maize and Wheat Improvement Center (CIMMYT) at Ciudad Obregón, México over four breeding cycles. We observed that single-kernel models using hyperspectral reflectance-derived relationship matrices performed similarly or superior to marker-and pedigree-based genomic selection models when predicting within and across environments. Multi-kernel models combining marker/pedigree information with hyperspectral reflectance phentoypes had the highest prediction accuracies; however, improvements in accuracy over marker-and pedigree-based models were marginal when correcting for days to heading. Our results demonstrates the potential of hyperspectral imaging in predicting grain yield within a multi-environment context, it also supports further studies on integration of hyperspectral reflectance phenotyping in breeding programs.

## INTRODUCTION

The aim of plant breeding programs is to develop and deliver new, high-yielding crop varieties that are adapted to a range of environmental conditions. Two major challenges of breeding for multiple environments are: (1) being able to account for the presence of genotype-by-environment interaction (*G* × *E*), and (2) the high costs of evaluating trials at multiple locations. Statistical, genomics, and phenomics tools that enable the accurate prediction and selection of candidate lines appropriate for each target environment may serve to increase the rate of genetic gain while reducing costs associated with large-scale multi-environment field trials.

In recent years, genomic selection (GS) and high-throughput phenotyping (HTP) have emerged as potential technologies for improving breeding efficiency (Furbank and Tester, 2011; Cabrera-Bosquet *et al.* 2012; Crossa *et al.* 2017). In GS, genome-wide marker effects are estimated for a “training set” of lines that has been phenotyped and genotyped (Meuwissen *et al.* 2001). Those estimates are then applied to selection candidates prior to phenotyping to predict trait values, which may reduce the time and cost of testing breeding lines. While GS was initially developed to predict within individual environments, recent studies have extended the genomic best linear unbiased prediction (GBLUP) model to accommodate *G* × *E* interactions. Burgueño *et al.* (2012), Heslot *et al.* (2013), JarquÍn *et al.* (2014), López-Cruz *et al.* (2015), and Pérez-RodrÍguez *et al*. (2017) reported increases in prediction accuracies with extended models relative to single-environment analyses.

HTP is based on the remote and proximal sensing of a large number of crop plants to collect relevant phenotypes while reducing labor time and cost (White *et al.* 2012; Araus and Cairns, 2014; Pauli *et al.* 2016). When deployed across different developmental growth stages and in multiple environments, HTP can drastically increase the phenotypic information available to breeding programs, which may help to improve selection accuracy. Although HTP traits represent indirect estimations and may not be able to provide the level of precision of direct measurement, they may be of particular use at the early generation stage in breeding programs when seed is limited. In wheat breeding, large numbers of lines are sown in small unreplicated plots for the purpose of visual selection and seed increase prior to grain yield testing in large replicated plots. Measurements of grain yield from small plots are not meaningful, nor are they feasible to collect when thousands of lines are being screened. Therefore, acquiring accurate predictions of grain yield at the early generation stage using HTP may serve to improve selection accuracy.

A range of ground-based and aerial HTP platforms have been recently developed to improve the accuracy, efficiency, and scope of phenotypic data collection (Andrade-Sanchez *et al.* 2014; Crain *et al.* 2016; Haghighattalab *et al.* 2016). A major advantage of aerial platforms is their ability to phenotype large areas of field trials in minimal time. This enhanced efficiency increases the spatial and temporal resolution of the phenotypic data and may be critical when assessing breeding trials at multiple locations. A number of recent studies have integrated traits collected with HTP into GS to increase prediction accuracy for grain yield in wheat (Rutkoski *et al.* 2014; Sun *et al.* 2017; Crain *et al.* 2018).

To date, many of the applications of aerial HTP within plant breeding have focused on measuring the spectral reflectance of the crop canopy. Plant cells, tissues, and pigments have wavelength-specific light absorption, reflectance, and transmittance patterns that may, for example, differentiate between healthy and stressed plants (Li *et al.* 2014). Vegetation indices (VIs) provide a convenient way of summarizing spectral reflectance information into scores that may be predictive of economically important traits (Govaerts *et al.* 1999). VIs such as the normalized difference vegetation index (NDVI) have been shown to be predictive of grain yield in wheat (Aparicio *et al.* 1999; Labus *et al.* 2002). However, since VIs are calculated from only a few wavelengths, they cannot leverage the high density of information captured by hyperspectral cameras, which record reflectance at a large number of narrowband wavelengths in the visible and near infrared regions of the light spectrum (Viña *et al.* 2011). While hyperspectral data may have a greater capacity than VIs to detect phenotypic differences between individuals, the high dimensionality of the data may complicate the interpretation. For these data to become meaningful for plant breeding, methods that can derive useful information on traits relevant to breeding are needed.

To address the high dimensionality of hyperspectral data, Aguate *et al.* (2017) found that integrating information from all hyperspectral wavelengths using ordinary least squares, partial least squares, and Bayesian shrinkage resulted in higher prediction accuracy than what could be achieved using individual VIs in maize. Montesinos-López *et al.* (2017a) proposed a Bayesian functional regression analysis using hyperspectral wavelengths that likewise resulted in higher accuracies for predicting grain yield in wheat when compared to a range of VIs. Montesinos-López *et al.* (2017b) further extended this method to incorporate genomic and pedigree information, in addition to accommodating *G* × *E* by modeling hyperspectral band-by-environment (*B* × *E*) interactions. Their study found that models that included the *B* × *E* term had higher prediction accuracies than those that did not, suggesting that hyperspectral reflectance may be a useful phenotype for modeling *G* × *E* interactions.

When collecting hyperspectral data within a multi-environment context, the number of predictors increases in proportion to the number of environments and phenotyping time-points observed, which may come at a computational cost depending on the type of prediction model used (Montesinos-López *et al.* 2017b). One possible approach that may minimize computation time would be to use the hyperspectral bands as a high dimensional predictor set, similar to the case of prediction with genomic markers in GBLUP, that is, to create a relationship matrix between individuals using the hyperspectral bands. This way, the number of bands could be very large without increasing the complexity of the GBLUP prediction model. Separate genomic marker/pedigree and hyperspectral reflectance kernels could be integrated to model the main genetic effects and *G* × *E* interactions, respectively.

Multi-environment field trials are often unbalanced, which can complicate their use in prediction across environments. When deploying HTP in large breeding programs, it can be difficult to ensure that all locations are phenotyped at the same stage and the same number of times throughout the season. Weather conditions, technical difficulties with the cameras or sensors, and scheduling with contracted pilots/airports may prohibit the use of aerial HTP on certain days, and lines grown at different sites may develop at faster or slower rates depending on weather and management conditions. As a result, sites may have different number of observed HTP time-points, as in Sun *et al.* (2017). It is well documented that canopy spectral reflectance varies according to crop phenology (Viña *et al.* 2003; Zhang *et al.* 2003). One strategy for comparing HTP time-points across varying sites may be to classify them according to the predominant developmental growth stage at the time of phenotyping.

To test these approaches, we deployed HTP on the CIMMYT Bread Wheat Improvement Program’s multi-environment yield evaluations of advanced germplasm to phenotype a range of differentially managed treatments with a hyperspectral camera at multiple time-points throughout the growing season. The main objectives of this research were to: (1) propose a multi-kernel, multi-environment GBLUP model that involves modeling genetic main effects using genomic markers or pedigrees and modeling *G* × *E* interactions using relationship matrices derived from hyperspectral reflectance data, (2) compare the prediction accuracies of models using genomic marker/pedigree main effects kernels and hyperspectral *G* × *E* interaction kernels separately and in combination, and (3) determine the optimal developmental growth stages for hyperspectral phenotyping based on grain yield prediction accuracy.

## MATERIALS AND METHODS

### Experimental Data

The dataset included a total of 3,771 bread wheat lines evaluated at the Campo Experimental Norman E. Borlaug in Ciudad Obregón, México over the course of four breeding cycles. In each cycle, lines were sown under five differentially managed treatments: Optimal Bed, Optimal Flat, Moderate Drought, Severe Drought, and Heat. Descriptions of the managed treatments are given in Table 1. The 20 managed treatment-breeding cycle combinations are herein referred to as site-years. Within each of the five managed treatments, 1092 lines were arranged into 39 trials in a α-lattice design with three replicates and six incomplete blocks per replicate. Each replicate contained two repeated checks. ‘Kachu #1’ was sown as a check in the Optimal Bed, Optimal Flat, and Moderate Drought managed treatments. ‘Baj #1’ was used as a check in the Severe Drought and Heat managed treatments. ‘Borlaug100 F2014’ was sown as the other check in all managed treatments. All lines within a breeding cycle were evaluated in all five managed treatments, while no lines other than checks overlapped between cycles. Records for some lines were removed for the analysis due to unavailability of genotypic or agronomic data, resulting in the final dataset: 588 lines in 2013-14, 1,033 lines in 2014-15, 1,063 lines in 2015-16, and 1,087 lines in 2016-17, for a total of 56,565 plots phenotyped. Each breeding cycle contained full-sib families with an average of two full-sibs per family.

**TABLE 1:** Description of the field management conditions in each of the five managed treatments sown at CIMMYT Research Station in Ciudad Obregon, México

| MANAGED<br>TREATMENT | PLANTING<br>DATE | PLOT<br>TYPE | PLOT<br>DIMENSIONS | IRRIGATION<br>METHODS |
| --- | --- | --- | --- | --- |
| Optimal Bed | Late November/<br>Early December | Two beds with<br>3 rows per bed | 2.8m × 0.8m | Five furrow irrigations |
| Optimal Flat | Late November/<br>Early December | Flat sown plot<br>with 6 rows | 4.0m × 1.3m | Five flood irrigations |
| Moderate<br>Drought | Late November/<br>Early December | Two beds with<br>3 rows per bed | 2.8m × 0.8m | Two furrow irrigations |
| Severe<br>Drought | Late November/<br>Early December | Flat sown plot<br>with 6 rows | 4.0m × 1.3m | Three minimal<br>irrigations through drip |
| Heat | Late February | Two beds with<br>3 rows per bed | 2.8m × 0.8m | Five furrow irrigations |

Grain yield (GY) in t ha^-1^ and lodging evaluated on an ordinal scale (0: no lodging; 5: completely lodged) were assessed in all three replicates in each site-year. Heading date was recorded as the date of 50% spike emergence within the plot. Maturity date was recorded as the date of senescence of the peduncle for 50% of stems in the plot. Days to heading (DTHD) and days to maturity (DTMT) were measured as the number of days to reach heading and maturity, starting from the date of the first irrigation if sowing was on dry soil or from the sowing date if sowing was in pre-irrigated fields. DTHD and DTMT were assessed for the first replicate only in each site-year due to the high heritability of the traits.

### Hyperspectral Phenotyping

Hyperspectral reflectance data were collected with a hyperspectral camera (A-series, Micro-Hyperspec VNIR, Headwall photonics Fitchburg, MA, USA) as part of the Alava Remote Sensing Spectral Solution (ARS3, Alava Ingenieros, Madrid, Spain) mounted in a Piper PA-16 Clipper aircraft. The camera’s sensor had a 12-bit radiometric resolution, covering the light spectrum in the 400–850nm region with a 7.5nm full width at half maximum and was set with an integration time of 18ms. The flights were scheduled around noon (GMT-7) and aligned to the solar azimuth angle at a height of 300m, resulting in 30cm resolution. The aerial imagery acquisition during the growing cycle was spaced approximately at seven-to ten-day intervals on mostly clear days.

Radiometric calibration of the sensor was done using coefficients derived from a calibrated uniform light source and an integrating sphere (CSTMUSS2000C Uniform Source System, LabSphere, North Sutton, NH, USA). Dark frame correction was performed for each flight dataset. Atmospheric calibration was performed using irradiance measurements acquired at the beginning and end of each flight using a Jaz spectrometer with a CC-3 Cosine Corrector (Ocean Optics Inc, FL, USA) for the 2016-17 breeding cycle. For remaining three breeding cycles, irradiance was modeled using aerosol optical depth from sun-photometer measurements (Microtops II, Solar Light Company, Glenside, PA, USA) based on the SMARTS simulation model (Gueymard, 1995, 2001).

Ortho-rectification and georeferencing of the imagery were performed using PARGE (ReSe Applications Schläpfer, Wil, Switzerland) based on data from the intertial navigation system (INS) attached to the camera (IG-500N model, SGB systems S. A. S., Carrières-sur-Seine, France). Hyperspectral reflectance data were extracted from the aerial images using the mean value of the pixels inside the central area of each observed plot, avoiding 0.5m from the plot border.

Each hyperspectral phenotyping time-point within each site-year was assigned a developmental growth stage classification according to the predominant growth stage of the lines within the site-year at the time of phenotyping (Table 2). The vegetative (VEG) stage was defined as the period between germination and 50% of plots at heading. The heading stage (HEAD) was defined as the period between 50% of plots at heading and 100% of plots at heading. The grain fill stage (GF) was defined as the period between 100% of plots at HD and 100% of plots at maturity.

**TABLE 2:**
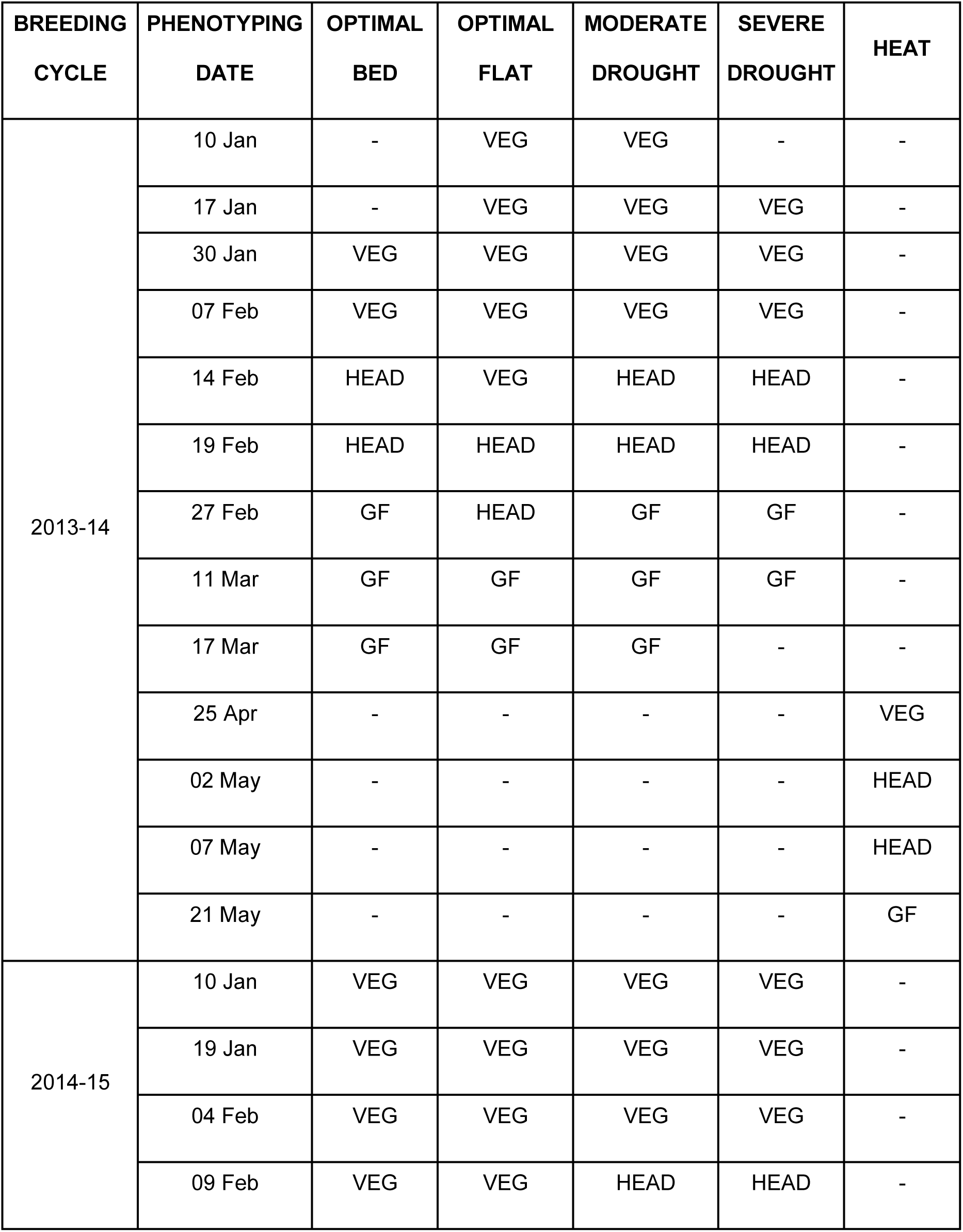
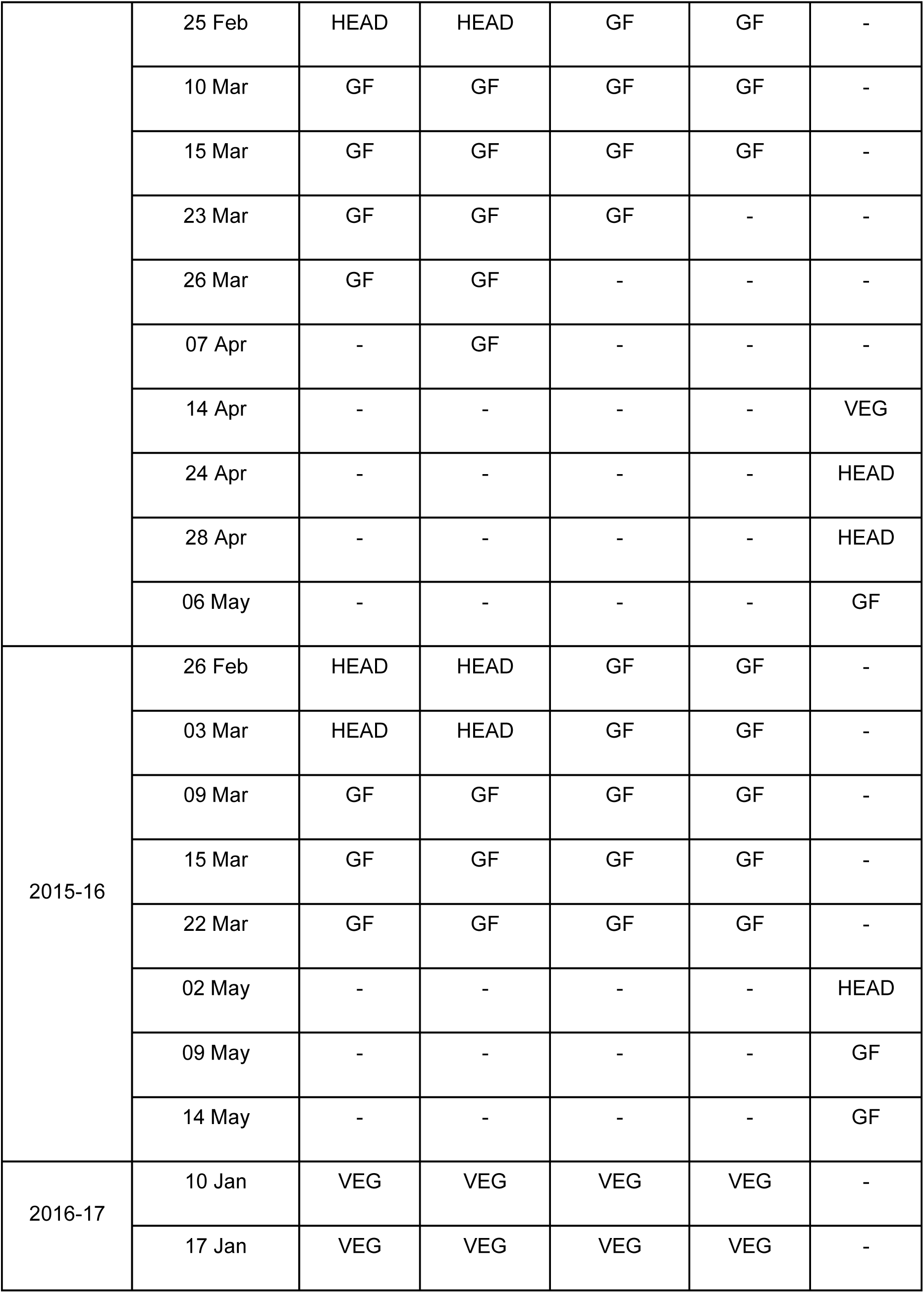
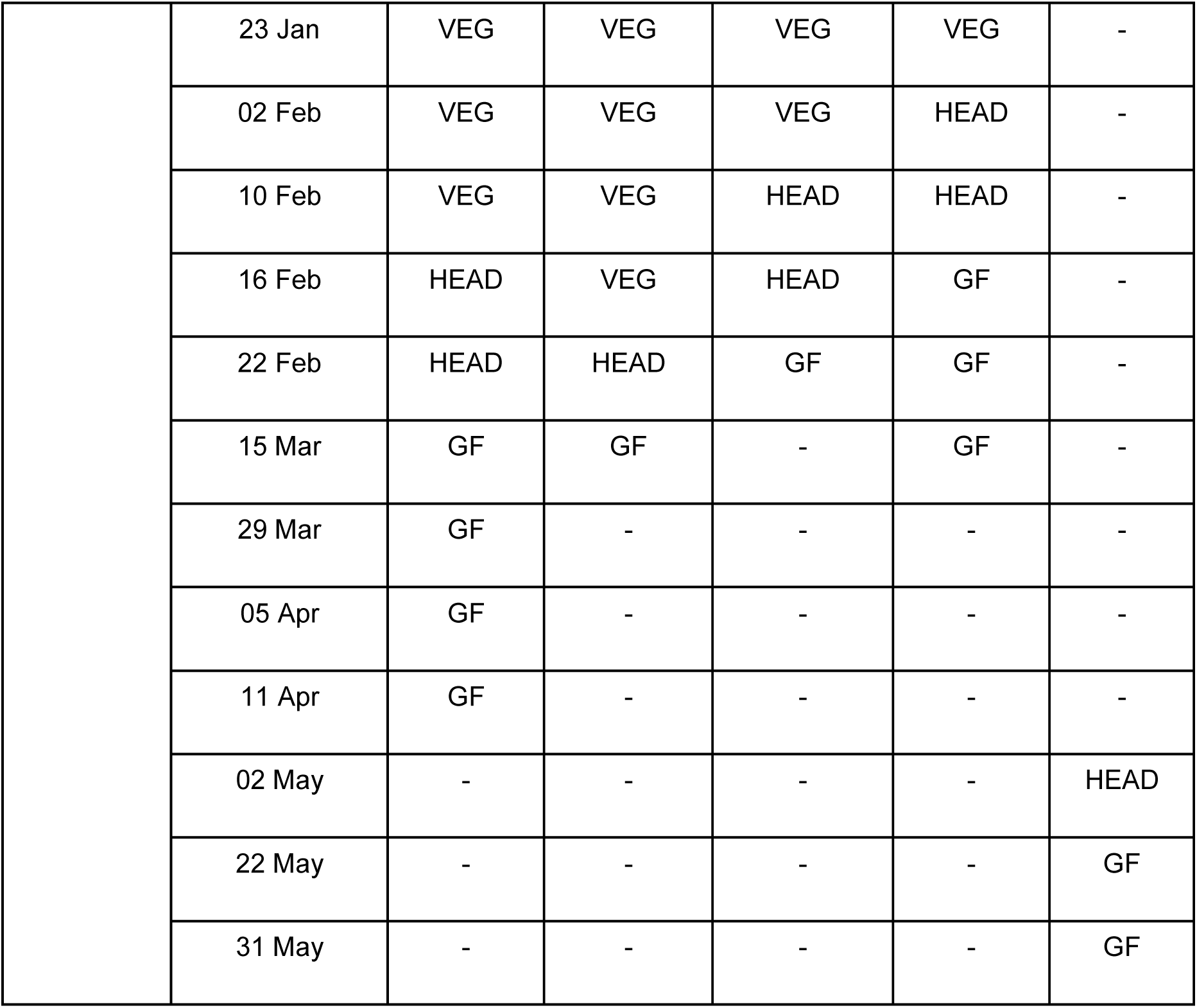
Classification of hyperspectral phenotyping time-points as vegetative (VEG), heading (HEAD), and grain fill (GF) phenological stages

### Genotypic Data

All lines were genotyped using genotyping-by-sequencing (Elshire *et al.* 2011) according to the pipeline described in Poland *et al.* (2012). From the initial set of 34,900 single nucleotide polymorphisms (SNPs), 8,519 SNPs remained after excluding all markers with more than 70% of missing data or minor allele frequency less than 0.05. For each marker, missing data were imputed using the sample mean of observed values (Poland *et al.*, 2012).

### Basic Statistical Models

Within each site-year, best linear unbiased estimates (BLUEs) were calculated for the agronomic traits and for each hyperspectral band at each phenotyping time-point using the following model:

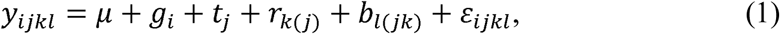

where *y*_*ijkl*_ is the trait value for genotype *i* within trial *j*, replicate *k*, and block *l*; *μ* is the overall mean; *g*_*i*_ is the fixed effect for genotype *i; t*_*j*_ is the random effect for trial *j*, which are assumed to be independently and identically distributed according to a normal distribution with mean zero and variance, σ_*t*_^2^that is, *t*_*j*_ ∼*iid N*(0, σ_*t*_^2^); *r*_*k(j)*_ ∼*iid N*(0, σ_*r*_^2^); is the random effect for replicate *k* within trial *j*; *b*_*l(jk)*_ ∼*iid N*(0, σ_*b*_^2^) is the random effect for block *l* within replicate *k* and trial *j*; and 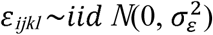 is the residual effect. For DTHD and DTMT, which were recorded for only one replicate, the replicate and block effects were excluded from the model. For validation of prediction models, best linear unbiased predictions (BLUPs) for GY within each site-year were calculated by treating the effect for genotype *g*_*i*_ in model (1) as random with *g*_*i*_ ∼*iid N*(0, σ_*g*_^2^) and by including a covariate for lodging as a fixed effect, fit only when lodging was observed within a site-year. To correct for the influence of phenology, BLUPs for GY were calculated again using DTHD included as a fixed effect in model (1).

BLUEs were also calculated for each hyperspectral band on a development growth stage basis according to the growth stage classifications detailed in Table 2. BLUEs for each hyperspectral band for the VEG, HEAD, and GF stages were estimated by fitting the following model:

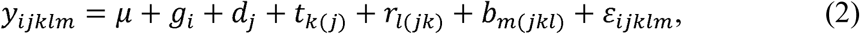

where *y*_*ijkl*_ is the hyperspectral reflectance trait value for genotype *i* on time-point *j* and within trial *k*, replicate *l*, and block *m*; *μ* is the overall mean, *g*_*i*_ is the fixed genetic effect for genotype *i*; *d*_*j*_ ∼*iid N*(0, σ_*d*_^2^) is a random effect for time-point *j*; *t* ∼*iid N*(0, σ_*d*_^2^) is random effect for trial *k* nested within time-point *j*; *r*_*l(jk)*_ ∼*iid N*(0, σ_*r*_^2^) is the random effect for replicate *l* nested within time-point *j* and trial k; *b*_*m(jkl)*_ ∼*iid N*(0, σ_*b*_^2^) is the random effect for block *m* nested within time-point *j*, trial k, and rep; and 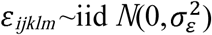 is the residual effect. BLUEs were also calculated for each hyperspectral band across all available phenotyping time-points (ALL) using model (2).

Broad-sense heritability within site-years was calculated for GY (Bernardo 2010) and each hyperspectral band as:

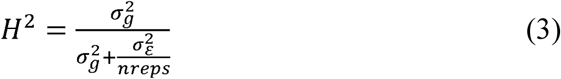

where σ *g*^2^ is the genetic variance, 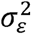 is the error variance, and *nreps* is the number of replicates (*nreps* = 3). For each breeding cycle, Pearson’s correlations for GY between managed treatments were calculated using the BLUEs derived from model (1). Pearson’s correlations were also calculated between GY BLUEs and the BLUEs for each hyperspectral band calculated in model (2).

### Relationship Matrices

Genetic relationships between individuals were modeled using genomic markers, pedigrees, and hyperspectral reflectance phenotypes. The genomic relationship matrix (***G***) was calculated according to Endelman and Jannink (2012). The additive relationship matrix (***A***) was derived from pedigrees and calculated as twice the coefficient of parentage.

Hyperspectral reflectance-based relationship matrices (***H***) were calculated within each site-year using the hyperspectral BLUEs calculated for 1) the individual time-points from model (1), 2) the developmental growth stages from model (2), and 3) all time-points from model (2). The matrices were calculated as 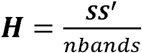, where ***S*** is a matrix of the centered and standardized BLUEs of the hyperspectral bands and *nbands* is the total number of hyperspectral bands observed.

### Prediction Models

#### Genetic Main Effects

To compare the utility of hyperspectral reflectance-based models to standard genomic prediction models, the following single-kernel genetic main effects model was fitted using marker-and pedigree-derived relationship matrices:

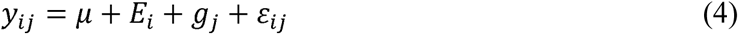

where *y*_*ij*_ is the BLUE of GY for genotype *j* in site-year *i, μ* is the overall mean, *E*_*i*_ is the fixed effect for site-year (*i=1,…,I), g*_*j*_ is the random effect for genotype *j* (*j=1,…,J*), and ε _*ij*_ is the residual effect. We assume that the joint distribution of genotype effects is distributed according to a multivariate normal distribution with mean **0** and variance-covariance matrix σ_*g*_^2^***K***, that is,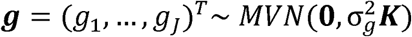, where 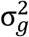 denotes the genomic variance and ***K*** represent the genomic relationship matrix between individuals ***G* (*K****=****G*)** or the numerical relationship matrix ***A*** (***K****=****A***).

#### Hyperspectral Reflectance Main Effects

To mimic a situation in which genomic marker and pedigree information are not available, the following single-kernel model was fitted using the hyperspectral reflectance-derived relationship matrices only:

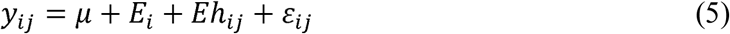

where *y*_*ij*_, *µ*, and *E*_*i*_ were as defined in model (4). *Eh*_*ij*_ is the random effect of the hyperspectral bands for genotype *j* in site-year i with the joint distribution of the hyperspectral bands as 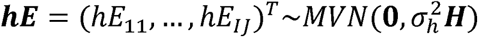, where 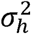 is the hyperspectral band variance and ***H*** is the hyperspectral reflectance-derived relationship matrix. Unlike matrices ***G*** and ***A***, which have only one row and column for each unique genotype evaluated, the matrix ***H*** has a row and column for each unique site-year-genotype combination where *I* × *J* results in the total number of site-year-genotype combinations.

#### Genetic Main Effects + Genetic G × E

As a basis for comparison to assess the advantage of *G* × *E* models that integrate both markers or pedigrees with hyperspectral reflectance phenotypes, model (4) was extended to accommodate *G* × *E* interactions using marker or pedigree information. The following multi-kernel model was fit:

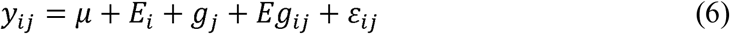

where *y*_*ij*_, *µ, E*_*i*_, and *g*_*j*_ were as defined in model (4). The *gE*_*ij*_ term was assumed to have multivariate normal distribution 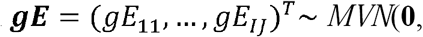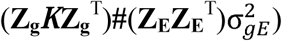 where **Z_g_** and **Z_E_** are incidence matrices for genotypes and site-years, ***K*** represents the genomic relationship matrix **(*K****=****G*)** or the numerical relationship matrix ***A*** (***K****=****A***), and 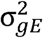 is the variance component for *gE*_*ij*_ (JarquÍn *et al.* 2014).

#### Genetic Main Effects + Hyperspectral Reflectance G × E

Finally, a multi-kernel model using marker or pedigree information to estimate the genetic main effects and hyperspectral reflectance phenotypes to model the *G* × *E* interactions was fitted:

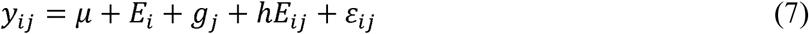

Here, *y*_*ij*_, *µ, E*_*i*_, and *g*_*j*_ were defined as above in model (4). The term *hE*_*ij*_ was assumed to have multivariate normal distribution 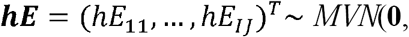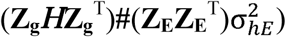 where **Z_g_** and **Z_E_** are incidence matrices for genotypes and site-years and 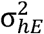 is the variance component for *hE*_*ij*_. The (**Z_g_*H*Z_g_**^T^)**#**(**Z_E_Z_E_**^T^) term was obtained with the block diagonal matrix *BDiag*(***H***_11_,…,***H***_II_) where ***H***_ii_ represent the hyperspectral relationship matrices for genotypes in site-year *i=1,…,I*.

### Software

Processing of the hyperspectral images was performed using the ARS3 hyproQ software (Álava Ingenieros, Madrid, Spain). Plot polygons for tabular data extraction were generated using ArcGIS (ESRI, Redlands, California, US). Images were aligned manually in ArcGIS if they did not overlay the plot polygons due to INS inaccuracy.

All models were fit using the R statistical programming language (R Core Team, 2018). Basic models (1-3) were fit with the package “ASReml-R” (Gilmour *et al.*, 2014) for R, while the prediction models (4-7) were fit using the “BGLR” package (de los Campos and Pérez-RodrÍguez, 2014) for R. The marker-based genetic relationship matrices were calculated using the A.mat() function within the “rrBLUP” package (Endelman 2011) for R. The coefficients of parentage were estimated using the “Browse” application within the International Crop Information System software package (McLaren *et al.* 2000).

### Assessing Model Prediction Accuracy

The above models were fit to assess model prediction accuracy for three prediction strategies representing different testing and evaluation problems relevant to plant breeding programs: 1) **within site-year**, 2) **within breeding cycle/across managed treatments**, and 3) **across breeding cycles/within managed treatment**.

A train-test (TRN-TST) validation scheme (Daetwyler *et al.*, 2013) of 20 random partitions was used to assess model prediction accuracy for all prediction strategies. For within site-year prediction, a random 80% of records within a given site-year were assigned to the TRN set, and the remaining 20% were used as the TST set for prediction. This reflects a scenario in which prediction of the trait values for a set of unobserved lines is performed within a managed treatment and breeding cycle of interest. Models (4) and (5) were fitted without the *E*_*i*_ term for site-year and models (6) and (7) were not tested for the within site-year prediction scheme because multiple site-years were not considered.

For prediction within breeding cycle/across managed treatments, predictions were carried out across the five managed treatments within a single breeding cycle. The TRN set consisted of all records from four out of the five managed treatments within a breeding cycle plus 20% of records from the fifth managed treatment. The remaining 80% of records from the fifth managed treatment were assigned to the TST set for prediction. In this situation, lines have been previously characterized in all managed treatments except one, where only 20% of records were available.

For prediction across breeding cycles/within managed treatment, predictions were performed across the four breeding cycles but within a single managed treatment. The TRN set contained all records from three out of the four breeding cycles for a given managed treatment plus 20% of records from the fourth breeding cycle. The remaining 80% of records from the fourth breeding cycle were assigned to the TST set for prediction. This scenario predicts performance of unobserved lines that are 1-3 breeding cycles removed from lines in the training set.

Table S1 illustrates examples of TRN-TST partitioning for the three prediction schemes. For all prediction strategies, model prediction accuracy was assessed as the Pearson’s correlation between predictions for GY and GY BLUPs with and without correction for DTHD calculated in model (1). Reported values are the mean and standard deviation of the 20 random TRN-TST partitions implemented. The same partitioning “splits” were used to assess all models to ensure fair comparisons.

### Data Availability

All phenotypic and genotypic data required to confirm the results presented in this study are available on CIMMYT Dataverse link: hdl:11529/10548109. The “GID” column denotes the unique identifiers for the genotypes. Supplementary file sets, tables and figures are available at FigShare.

## RESULTS

### Descriptive Statistics

To evaluate the potential of integrating aerial hyperspectral reflectance phenotypes into GS to improve prediction accuracy for GY in wheat, we deployed an airplane equipped with a hyperspectral camera to phenotype wheat breeding trials in 20 site-years at multiple time-points throughout the growing season. Across the site-years, the Optimal Bed and Optimal Flat managed treatments had higher GY than the stressed Moderate Drought, Severe Drought, and Heat managed treatments (Figure 1). The standard deviations for GY remained relatively stable across site-years, ranging from 0.30 to 0.68 t ha^-1^ (Table S2). Broad-sense heritability for GY was high across site-years, ranging from 0.58 to 0.94 (Table S2). Overall, correlations for GY between managed treatments were moderately positive with the managed treatments that received similar levels of irrigation (e.g., Optimal Bed with Optimal Flat; Moderate Drought with Severe Drought) showing the strongest correlations (Figure S1).

**FIGURE 1:**
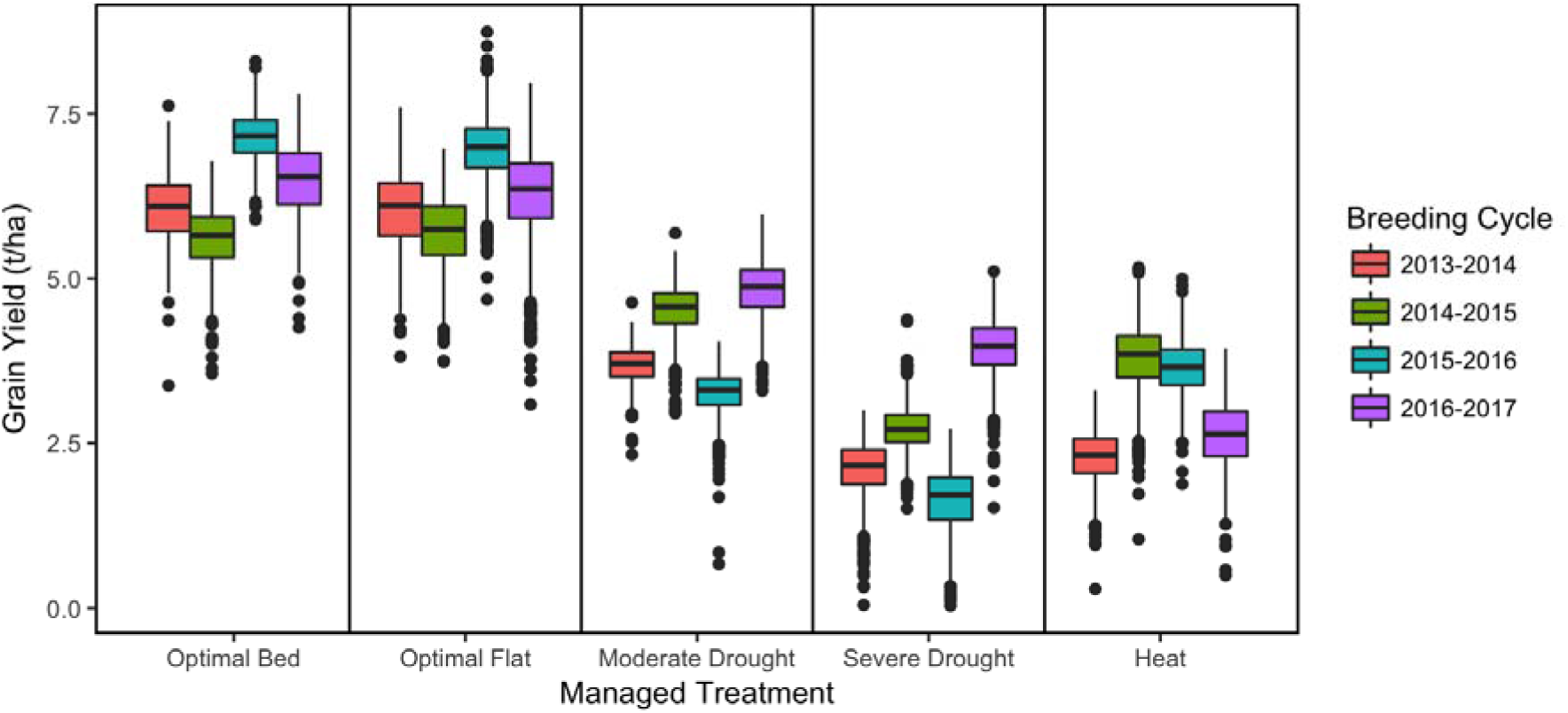
Boxplot of BLUEs for wheat grain yield (t ha^-1^) in each of the 20 observed site-years

Between 3 and 11 hyperspectral phenotyping time-points were collected within each site year (Table 2). In most site-years, at least one hyperspectral phenotyping time-point was collected during each of the three developmental growth stages. While the managed treatments within a breeding cycle were often phenotyped on the same date, the growth stage at the time of phenotyping often differed among the managed treatments (Figure 2). During the 2015-16 cycle, technological issues with the camera prevented early season phenotyping. As a result, no hyperspectral data were collected during the VEG stage. The Optimal Bed, Optimal Flat, and Heat managed treatments had hyperspectral phenotypes for the HEAD and GF stages only, while the Moderate Drought and Severe Drought managed treatments had hyperspectral phenotypes for the GF stage only due to early maturation of the crop under stress.

**FIGURE 2:**
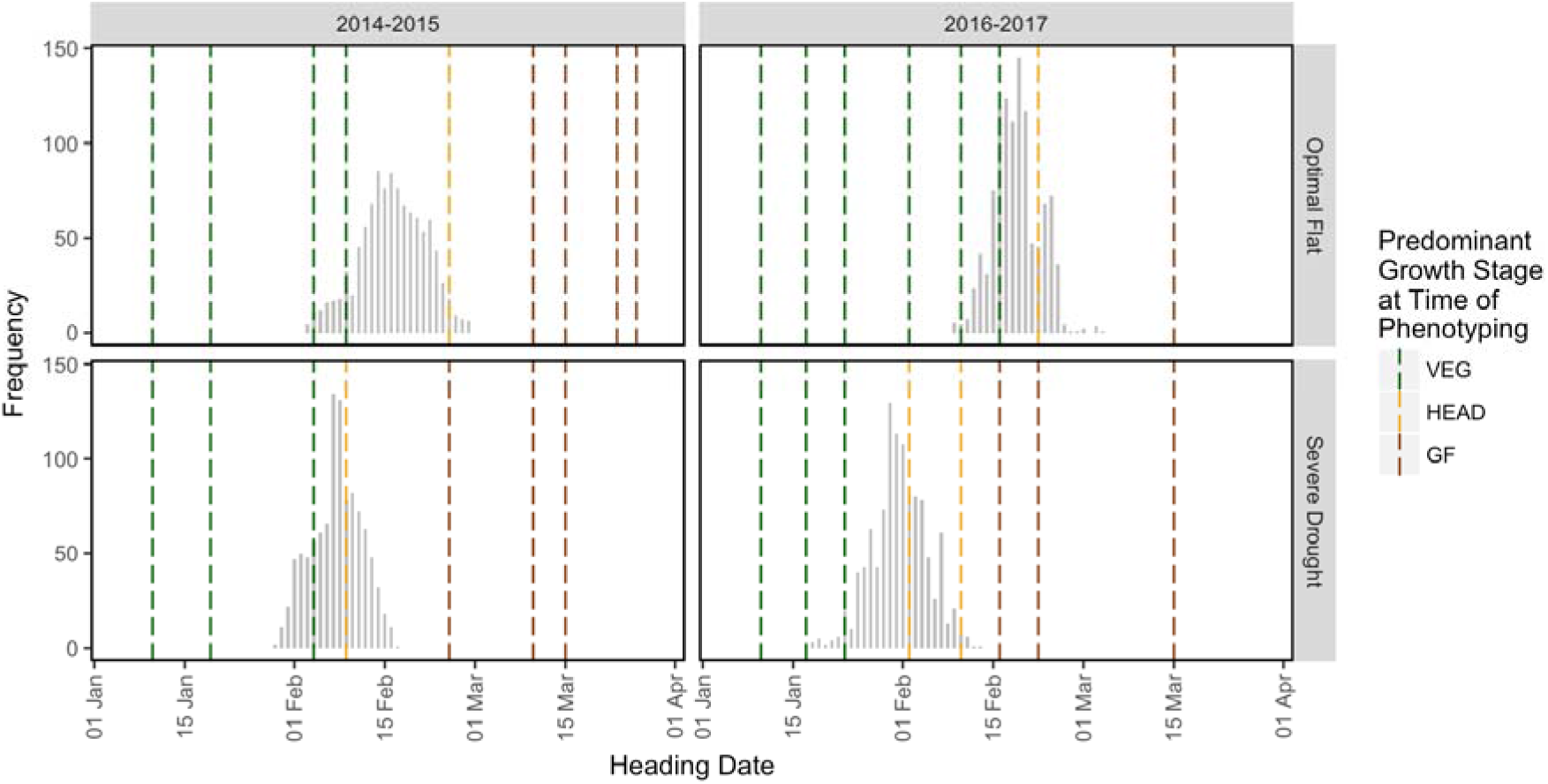
A graphical representation of the unbalanced nature of the hyperspectral reflectance phenotypic data Four site-years are represented: 2014-15 Optimal Flat, 2014-15 Severe Drought, 2016-17 Optimal Flat and 2016-17 Severe Drought. The histograms represent heading dates. Each dashed line corresponds to a hyperspectral phenotyping date colored according to the predominant growth stage of the lines at the time of phenotyping.

Broad-sense heritability estimates for the hyperspectral bands were generally between 0.5 and 0.8 for most phenotyping time-points (Figure 3). Heritabilities were relatively homogeneous across individual time-points and developmental growth stages. In all site-years, the lowest heritabilities were observed for the hyperspectral bands between 398 and 425 nm. Heritabilities in the 2013-14 Optimal Flat environment were notably lower than in other site-years, falling between 0 and 0.25, though the GF time-points were more heritable with values around 0.4. Low heritabilities (<0.2) were also observed for the 9 March time-point in the 2015-16 Moderate Drought environment.

**FIGURE 3:**
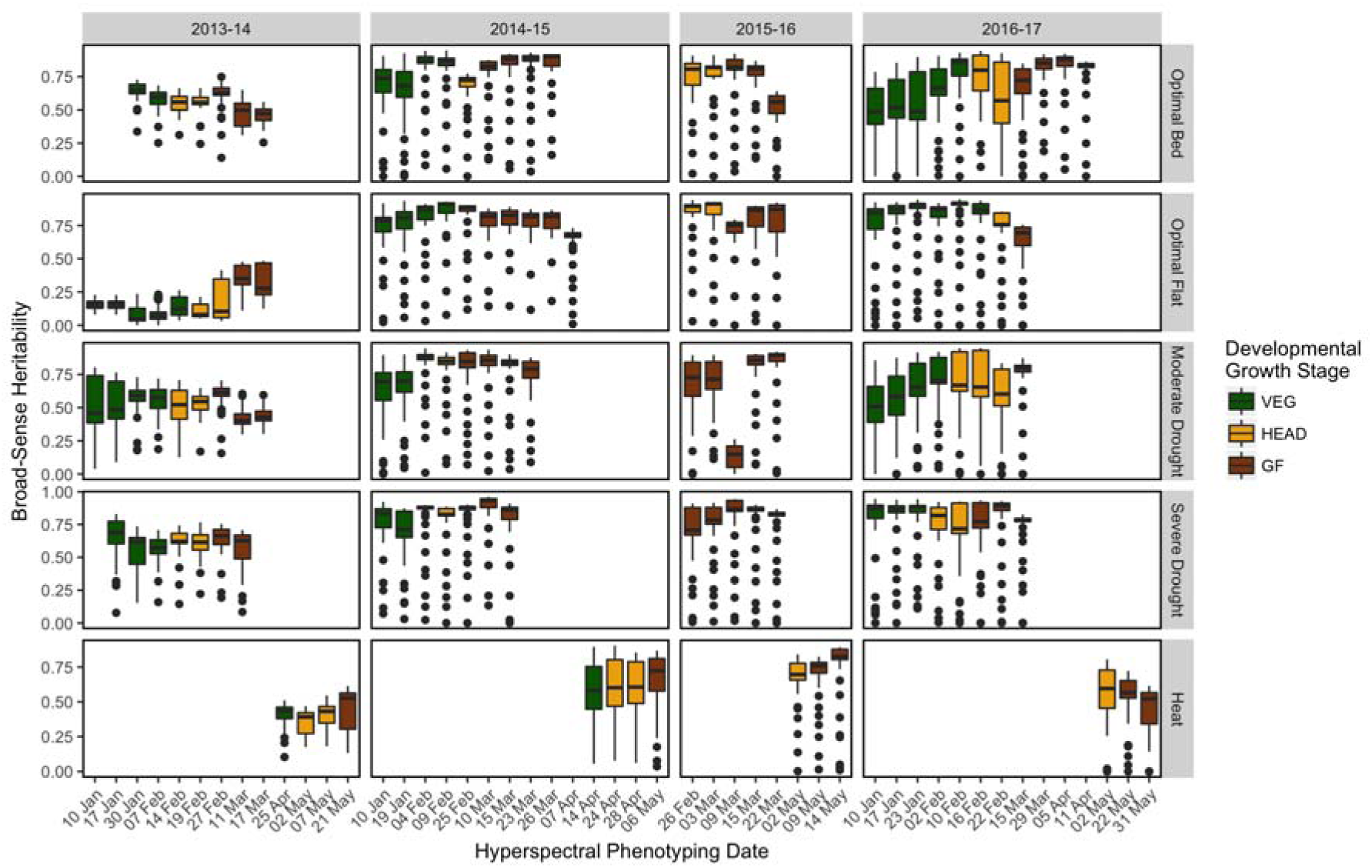
Broad-sense heritabilities of the hyperspectral wavelengths for each phenotyping time-point within each site-year Each boxplot represents the distribution of broad-sense heritability values for the 62 hyperspectral wavelengths observed. The colors correspond to the developmental growth stage of the site-year at the time of hyperspectral phenotyping.

Correlations between the individual hyperspectral bands and GY ranged from −0.68 to 0.64 across the 20 site-years and were most frequently the strongest above 500 nm (Figure 4). While there were some observable similarities in correlation patterns among time-points taken during the same growth stage within the same site-year, patterns were not uniform across site-years, breeding cycles, or managed treatments. In some site-years, there were clear differences in correlation patterns between developmental growth stages. For example, in the 2014-15 Optimal Bed site-year, correlations between GY and hyperspectral reflectance in the 398-700 nm range were around 0 during the VEG stage but become progressively more negative during the HEAD and GF stages, reaching around −0.4. For the same managed treatment during the 2016-17 breeding cycle, distinct differences in correlation patterns between the three developmental growth stages were also observed; however, the patterns do not reflect those observed in 2014-15. Despite irregularities in correlation patterns, the correlations between hyperspectral bands from the GF stages and GY were, on average, stronger by 0.10 than for hyperspectral phenotypes taken during the VEG and HEAD stages.

**FIGURE 4:**
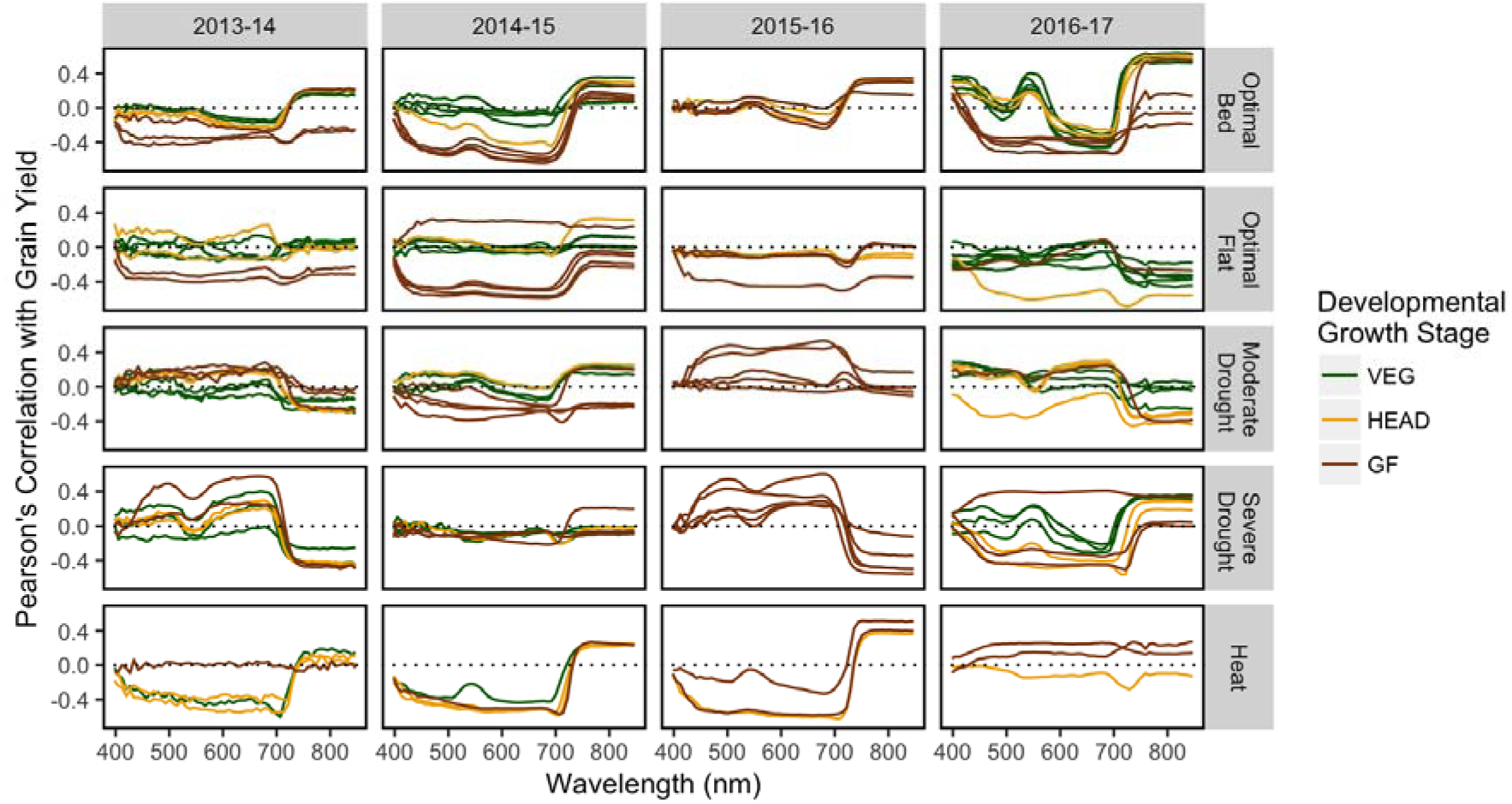
Empirical correlations between grain yield and hyperspectral reflectance BLUEs within each site-year Each line represents a phenotyping time-point and the correlation between grain yield and the 62 hyperspectral wavelengths observed. Lines are colored according to the predominant developmental growth stage of the site-year at the time of hyperspectral phenotyping.

### Model Prediction Accuracy

#### Within Site-Year

Five types of relationship matrices were used within models (4) and (5) for within site-year prediction: genomic (***G***), pedigree (***A***), individual hyperspectral phenotyping time-points (e.g. ***H*_*10Jan*_**, ***H*_*23Mar*_**, etc.), hyperspectral phenotype BLUEs calculated from multiple time-points for each developmental growth stage (***H*_*VEG*_**, ***H*_*HEAD*_**, ***H*_*GF*_**), and hyperspectral phenotype BLUEs calculated using all available time-points (***H*_*ALL*_**). Model (4) was used to assess accuracies using the ***G*** and ***A*** matrices for prediction. Model (5) was fit to assess accuracies using the hyperspectral relationship matrices.

Prediction accuracies of models using the individual hyperspectral time-point-derived relationship matrices ranged from 0.00 to 0.75 with a mean of 0.42 (Figure 5A). These results are similar to the prediction accuracies observed for ***G*** and ***A***, which recorded means of 0.41 and 0.42 and ranges of 0.19-0.60 and 0.19-0.67, respectively. In 18 and 17 of the 20 site-years, at least one time-point showed accuracies that were equivalent or superior to ***G*** and ***A***, respectively. In considering the most highly predictive time-point in each site-year, 11 were recorded during the GF stage, followed by 6 and 3 in the HEAD and VEG stages, respectively, although the site-years had different numbers of hyperspectral phenotyping time-points from each growth stage and some site-years had no observations recorded during the VEG and HEAD stages.

**FIGURE 5:**
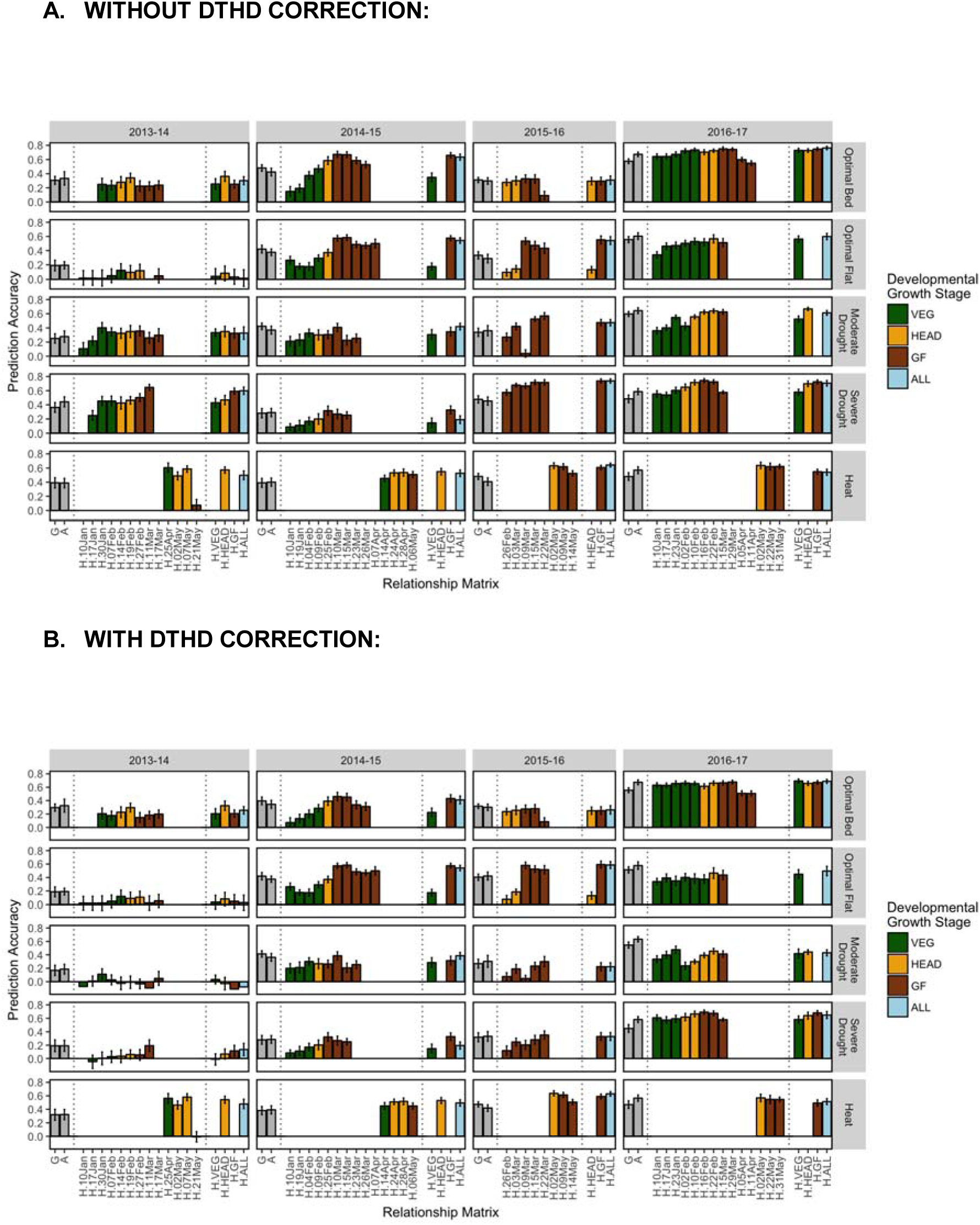
Within site-year prediction accuracies, with and without correction for DTHD Accuracy is expressed as the average Pearson’s correlation between predictions and observed BLUPs for GY across 20 random TRN-TST partitions. Results shown according to the type of relationship matrix tested: Genomic (**G)**, pedigree (**A**), individual hyperspectral time-points (e.g. **H10Jan**, **H.23Mar**, etc.), hyperspectral BLUEs for each developmental growth stage (**H.VEG**, **EAD**, **H.GF**), and hyperspectral BLUEs across all time-points (**H.ALL**). The color corresponds to the predominant developmental growth stage of the site-year at the time of phenotyping. Error bars are the standard deviation of prediction accuracy for the 20 random partitions

The level of prediction accuracy for the individual hyperspectral time-points was highly correlated with the strength of the relationship between hyperspectral reflectance and GY (Figure 6). For a given time-point within a site-year, the correlation between hyperspectral reflectance and GY was taken for each of the 62 wavelengths. The average of the absolute values of those correlations, or their relative magnitudes, was then compared to the level of prediction accuracy for that time-point. For the 71 time-point/site-year combinations, the average magnitude of the correlation between hyperspectral reflectance and GY explained 51 percent of the variation in prediction accuracy, though the two characters exhibited a non-linear relationship (Figure 6A). Correlations beyond 0.25 between hyperspectral reflectance and GY provided little marginal increase in prediction accuracy. When GY was corrected for DTHD, this relationship was linear and the correlation between reflectance and GY explained 26 percent of the variation in prediction accuracy (Figure 6B).

**FIGURE 6:**
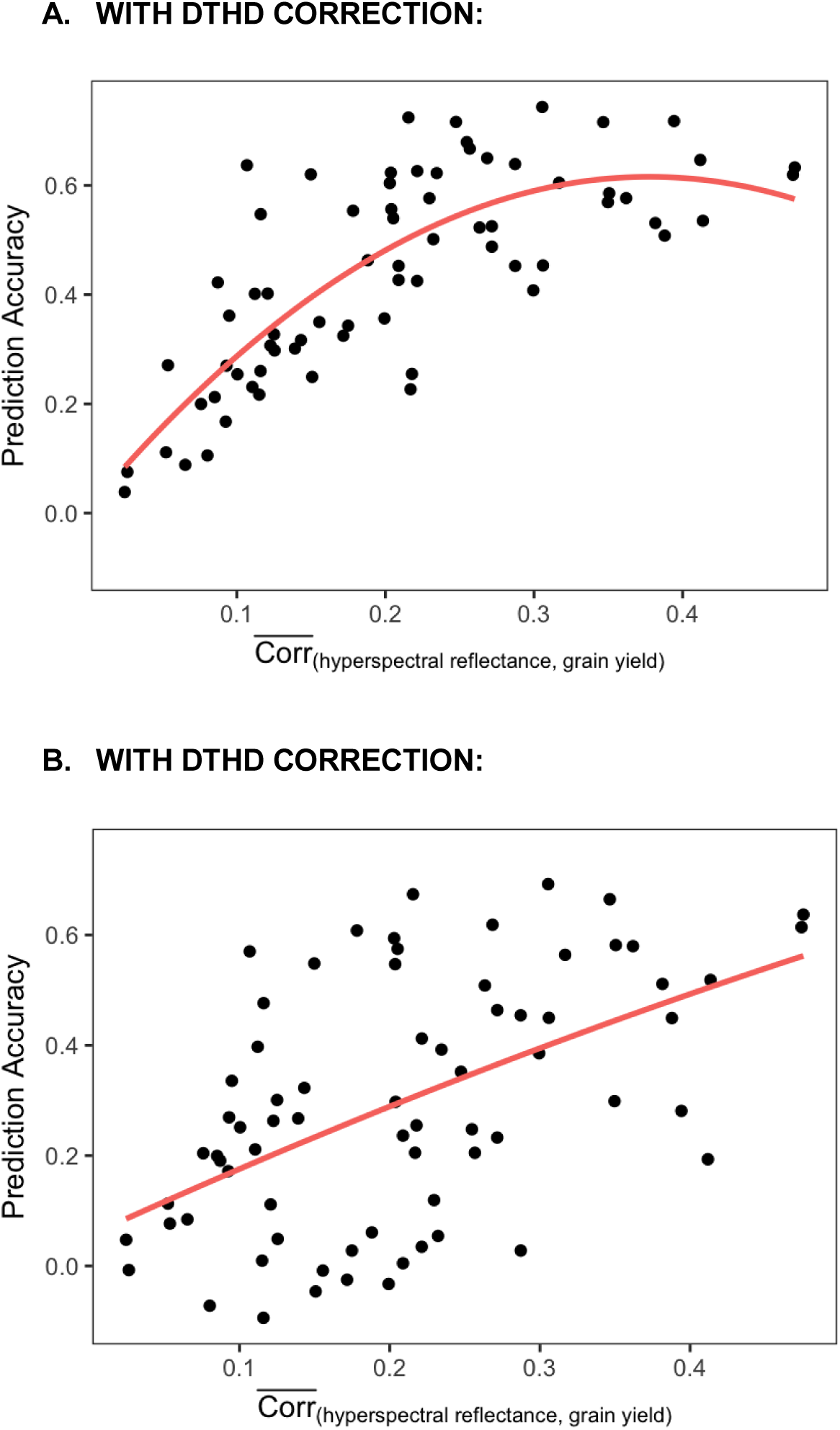
The relationship between individual hyperspectral time-point prediction accuracy and the average magnitude of the correlations between hyperspectral bands and GY The absolute values of the correlations between hyperspectral bands and GY were calculated and then averaged across the 62 bands for each time-point within each site-year. Plotted on the axis, this represents the average strength of the relationship between hyperspectral reflectance and GY for each time-point within each site-year. The y-axis shows the prediction accuracy for each individual hyperspectral time-point in within site-year prediction.

In considering accuracies using the hyperspectral phenotype BLUEs for each growth stage, ***H*_*GF*_** had the highest prediction accuracies in 11 of the 20 site-years, followed by ***H*_*HEAD*_** in 8 site-years and ***H*_*VEG*_** in 1 site-year. Combining multiple time-points did not necessarily increase accuracy. The accuracies of ***H*_*VEG*_**, ***H*_*HEAD*_**, and ***H*_*GF*_** were comparable to the mean of the accuracies of the individual time-points. Likewise, the accuracy of ***H*_*ALL*_**, which combined all available time-points, was on average slightly higher than the mean of the accuracies of the individual time-points but lower than the most predictive time-point.

While correcting for DTHD had negligible impacts on prediction accuracy with respect to the ***G*** and ***A*** models, the accuracies of the hyperspectral reflectance models decreased by 0.10 on average (Figure 5B). The greatest reductions were observed in the Severe Drought and Moderate Drought treatments, which showed average decreases in accuracy of 0.20 and 0.18, respectively. After the DTHD correction, at least one-time point recorded prediction accuracies that were equivalent or superior to the accuracies of ***G*** and ***A*** in 13 out of the 20 site-years,

#### Within Breeding Cycle/Across Managed Treatments

For prediction across managed treatments within a breeding cycle, the genomic, pedigree, and hyperspectral reflectance matrices were tested individually in single-kernel models and in combination in multi-kernel models (Figure 7). Single-kernel models included the following: genetic main effects (***G*** and ***A***) from model (4) and hyperspectral reflectance main effects (***H*_*VEG*_**, ***H*_*HEAD*_**, ***H*_*GF*_**, and ***H*_*ALL*_**) from model (5). Multi-kernel models were built by combining a main effects kernel with a *G* × *E* interaction kernel in models (6) and (7). In model (6), the *G* × *E* interaction kernel was also fit using ***G*** or ***A***. These models are herein referred to as ***G G***x***E*** and ***A G***x***E***. In model (7), the hyperspectral reflectance matrices were used to estimate the *G* × *E* interaction kernel. These models are referred to as ***G + H*_*VEG*_**, ***A + H*_*HEAD*_**, etc. according to the respective relationship matrices used to estimate the main and *G* × *E* interaction effects. The ***H*_*VEG*_**, ***G + H*_*VEG*_**, and ***A + H*_*VEG*_** models were assessed for 14 out of the 20 site-years due to the unavailability of hyperspectral data at the vegetative stage in the remaining 6 site-years. Likewise, the ***H*_*HEAD*_**, ***G + H*_*HEAD*_**, and ***A + H*_*HEAD*_** models were assessed for 18 out of the 20 site-years. The remaining models with ***H*_*GF*_** and ***H*_*ALL*_** were tested for all site-years.

**FIGURE 7:**
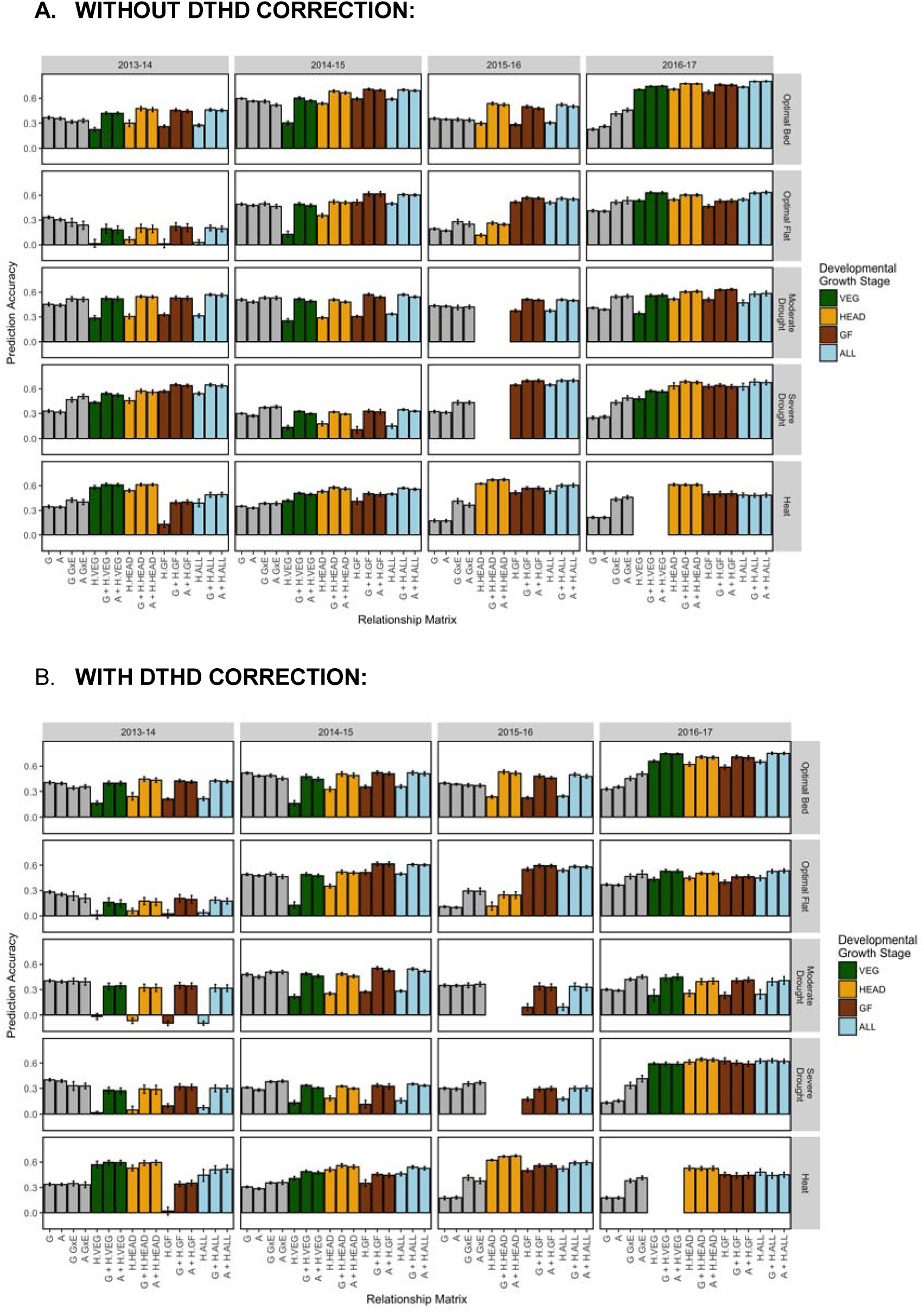
Within breeding cycle/across managed treatments prediction accuracies, with and without correction for DTHD Accuracy is expressed as the average Pearson’s correlation between predictions and observed BLUPs for GY across 20 random TRN-TST partitions. In each partition, the TRN set consisted of all records from four out of the five managed treatments within the breeding cycle plus 20% of records from the fifth managed treatment. Single-kernel models tested were genetic main effects only (**G** or **A**) and hyperspectral reflectance main effects plus GxE (**H.VEG, H.HEAD, H.GF, LL**). Multi-kernel models assessed were genetic main effects plus genetic GxE (**G GxE**, **A GxE**) and genetic main effects plus hyperspectral reflectance GxE (e.g., **G + H.VEG**, **A + H.ALL**, etc.). The color corresponds to the developmental growth stage of the site-year at the time of phenotyping. Error bars are the standard deviation of prediction accuracy for the 20 random partitions

When considering the single-kernel hyperspectral main effect models, 2013-14 Optimal Flat had prediction accuracies close to zero. The 2013-14 Optimal Flat site-year had high levels of lodging. This may have affected the reflectance signatures of the crop canopy. The hyperspectral reflectance phenotypes for 2013-14 Optimal Flat had lower heritabilities than all other site-years. To summarize trends for the remaining 19 site-years, the results from the single-kernel hyperspectral main effects models for 2013-14 Optimal Flat were removed as outliers. The accuracies of ***H*_*VEG*_** averaged 0.37 across the site-years where it was tested, which was similar to the accuracies of ***G*** (0.35) and ***A*** (0.34) (Figure 7A). The ***H*_*HEAD*_**, ***H*_*GF*_**, and ***H*_*ALL*_** showed slightly higher averages accuracies of 0.44, 0.44, and 0.46, respectively. The greatest differences in prediction accuracy between the hyperspectral reflectance-based models and the ***G*** and ***A*** models were observed in the 2015-16 and 2016-17 breeding cycles. In these cycles, the ***H*_*HEAD*_**, ***H*_*GF*_**, and ***H*_*ALL*_** models showed accuracies that were on average 0.22 greater than the **G** and **A** models. When correcting all models for DTHD, mean accuracies of the ***H*_*VEG*_** (0.29), ***H*_*HEAD*_** (0.34), ***H*_*GF*_** (0.30), and ***H*_*ALL*_** (0.34) models were more similar to the ***G*** (0.33) and ***A*** (0.32) models (Figure 7B).

Expanding the ***G*** and ***A*** models to account for the *G* × *E* interactions in the ***G G***x***E*** and ***A G***x***E*** improved prediction accuracies to a level of 0.43 on average, an increase of 0.08 and 0.09 over the ***G*** and ***A*** models, respectively. These improvements in accuracy were more pronounced during the 2016-17 breeding cycle, recording gains in accuracy of 0.16 and 0.19 for ***G G***x***E*** and ***A G***x***E***, respectively. Likewise, slightly greater improvements were observed for the Severe Drought (gains of 0.13 and 0.16 for ***G G***x***E*** and ***A G***x***E***, respectively) managed treatment. Similar trends were observed when correcting for DTHD, though accuracies were 0.04 lower on average.

The multi-kernel models that estimated the main effects using ***G*** or ***A*** and the *G* × *E* interactions using hyperspectral reflectance gave the highest accuracies overall. Hyperspectral reflectance matrices ***H*_*VEG*_**, ***H*_*HEAD*_**, ***H*_*GF*_**, and ***H*_*ALL*_** had average accuracies of 0.54, 0.56, 0.56, and 0.58, respectively, when integrated with ***G*** and average accuracies of 0.53, 0.55, 0.55, and 0.57, respectively, when integrated with ***A***. In most site-years, no clear “best model” could be identified among the multi-kernel models within a site-year. Accuracies were similar regardless of during which growth stage the hyperspectral data were recorded or whether genomic markers or pedigrees were used to model the main effects.

When compared to the ***G G***x***E*** and ***A G***x***E*** models, the use of hyperspectral reflectance to estimate the *G* × *E* interactions increased prediction accuracies by an average of 0.10 and 0.12, respectively, and by an average of 0.06 and 0.08, respectively after correcting for DTHD. Compared to the corresponding single-kernel hyperspectral reflectance models, the integration of markers or pedigree with hyperspectral reflectance in multi-kernel models increased prediction accuracies on an average by 0.11 and 0.17 respectively, and 0.14 and 0.19, respectively when correcting for DTHD.

#### Across Breeding Cycles/Within Managed Treatment

The models tested for prediction across breeding cycles/within managed treatment were the same as those tested for prediction within breeding cycle/across managed treatments (Figure 8). On average, accuracies were 0.08 lower in this prediction scheme than for prediction within breeding a cycle and across managed treatments.

**FIGURE 8:**
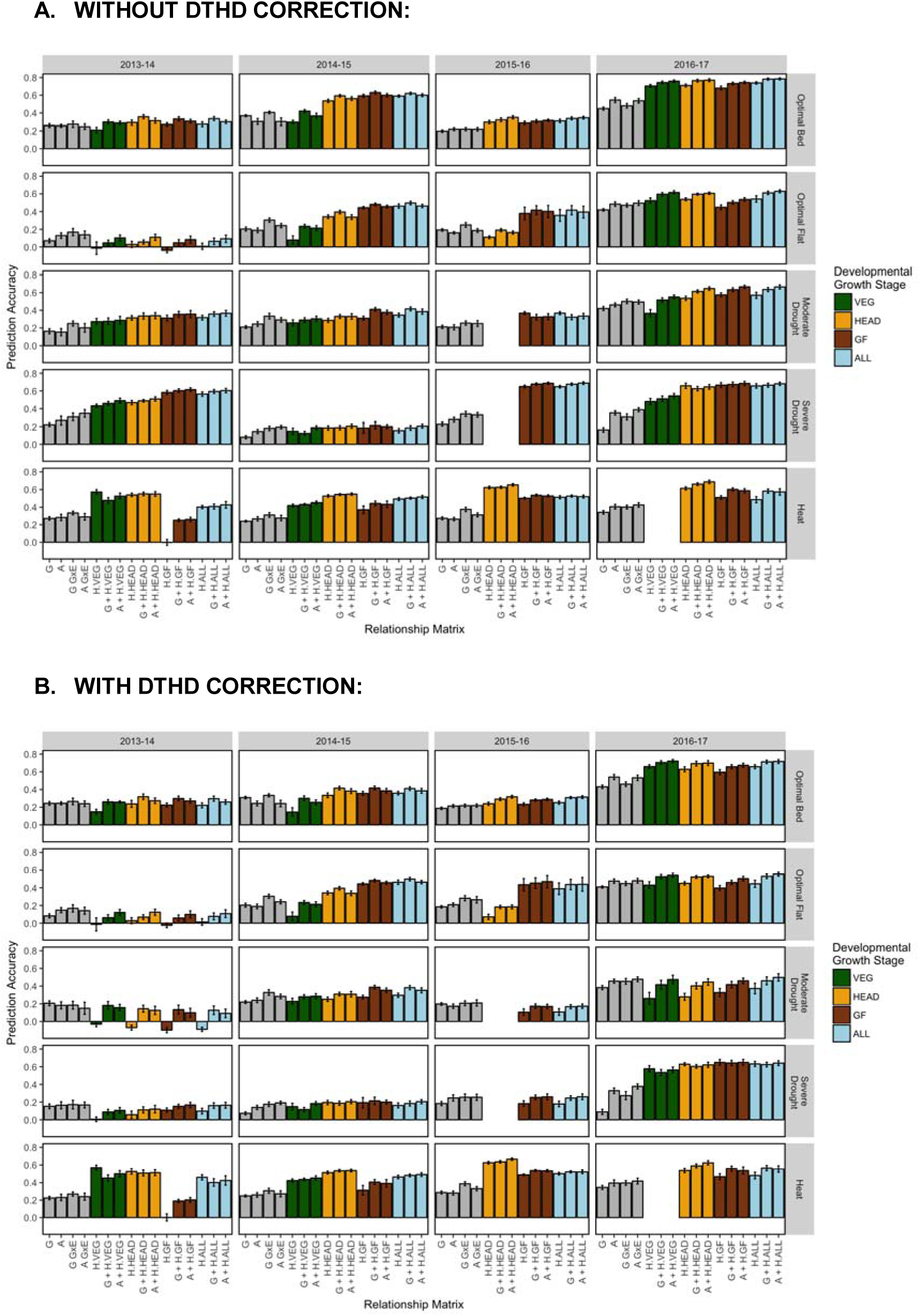
Across breeding cycles/within managed treatment prediction accuracies, with and without correction for DTHD Accuracy is expressed as the average Pearson’s correlation between predictions and observed BLUPs for GY across 20 random TRN-TST partitions. Single-kernel models tested were genetic main effects only (**G** or **A**) and hyperspectral reflectance main effects plus GxE (**H.VEG, H.HEAD, H.GF, H.ALL**). Multi-kernel models assessed were genetic main effects plus genetic GxE (**G GxE**, **A GxE**) and genetic main effects plus hyperspectral reflectance GxE (e.g., **G + H.VEG**, **A + H.ALL**, etc.). The color corresponds to the developmental growth stage of the site-year at the time of phenotyping. Error bars are the standard deviation of prediction accuracy for the 20 random partitions.

For the single-kernel hyperspectral main effects models, ***H*_*GF*_** in 2013-14 Heat and all single-kernel ***H*** models for 2013-14 Optimal Flat showed prediction accuracies close to zero. The 2013-14 Heat site-year was phenotyped once during the GF late in the growing season (21 May) which was close to maturity. Figure 4 shows that the correlation between hyperspectral reflectance and GY for that time-point is close to zero for each of the 62 wavelengths. To consider average trends, these were removed as outliers. Mean accuracies of the ***H*_*VEG*_**, ***H*_*HEAD*_**, ***H*_*GF*_**, and ***H*_*ALL*_** models were 0.37, 0.45, 0.45, and 0.46, respectively (Figure 8A). The accuracies of the ***G*** and ***A*** models were slightly lower, recording averages of 0.25 and 0.28, respectively. When correcting for DTHD, the mean accuracies of ***H*_*VEG*_**, ***H*_*HEAD*_**, ***H*_*GF*_**, and ***H*_*ALL*_** models were 0.28, 0.34,, and 0.34, respectively (Figure 8B). While closer in performance to ***G*** and ***A***, the single-kernel hyperspectral main effects models still conferred an advantage in some site-years.

Expanding the ***G*** and ***A*** models to account for the *G* × *E* interactions marginally increased accuracies to 0.32 and 0.31 for the ***G G***x***E*** and ***A G***x***E*** models, respectively. As with prediction within breeding cycles/across managed treatments, the highest accuracies were observed for the multi-kernel models that estimated the main effects using markers or pedigrees and the *G* × *E* interactions using hyperspectral reflectance. Hyperspectral reflectance matrices ***H*_*VEG*_** had an average accuracy of 0.39 when integrated with ***G*** and 0.41 when integrated with ***A***. ***H*_*HEAD*_**, ***H*_*GF*_**, and ***H*_*ALL*_** had average accuracies of 0.46, 0.46, and 0.48, respectively, when integrated with ***G*** or ***A***. As before, for most site-years, these multi-kernel models were observed to have similar levels of accuracy within each site-year, irrespective of which hyperspectral reflectance matrix was used to model the *G* × *E* interactions or whether genomic markers or pedigrees were used to model the main effects.

When compared to the ***G G***x***E*** and ***A G***x***E*** models, estimating the *G* × *E* interactions using hyperspectral reflectance increased prediction accuracy by an average of 0.12 and 0.14, respectively, and by an average of 0.07 and 0.08, respectively, when correcting for DTHD. Average improvements over the corresponding single-kernel hyperspectral reflectance main effects models ranged from 0.0.04 to 0.06 and from 0.06 to 0.08 after correcting for DTHD.

Mean accuracies and standard deviations of all models for the three prediction schemes can be found in Files S1, S2, and S3 respectively.

## DISCUSSION

We proposed a multi-kernel, across-environment GBLUP model that uses relationship matrices derived from genomic markers, pedigrees, and aerial hyperspectral reflectance phenotypes to estimate both genetic main effects and G × E interactions in the context of a wheat breeding program. Our study found that deriving a relationship matrix from high-dimensional hyperspectral reflectance phenotypes - as if they were genetic markers - for use in GBLUP can be an effective approach for predicting GY in wheat and in many situations resulted in predictions accuracies that were equivalent or superior to the use of genomic markers or pedigrees. This is consistent with similar studies that used linear and non-linear modeling approaches including ordinary least squares, partial least squares, Bayesian shrinkage, and functional regression methods to predict grain in maize and wheat with hyperspectral reflectance data (Aguate *et al.* 2017; Montesinos-Lopéz *et al.* 2017a).

The heritabilities of the hyperspectral wavelengths and their correlations with GY were not homogeneous across time-points, growth stages, breeding cycles or managed treatments. Wavelengths between 398 and 425 nm were observed to have the lowest heritabilities while those below 500 nm tended to have the weakest correlations with GY. These results are consistent with a similar study of canopy reflectance in wheat (Hansen *et al.* 2003) that found low signal-to-noise between 400 and 438 nm. According to Mahlein *et al.* (2015), the hyperspectral imaging systems most commonly used in agriculture have poor sensitivity to reflectance in the blue region of the light spectrum (400-500 nm). Despite this, Montesinos-Lopéz *et al.* (2017a) showed that removing wavelengths with low heritability did not improve prediction accuracies when using hyperspectral reflectance to predict GY in wheat. Based on this result, we used all hyperspectral wavelengths to build relationship matrices for prediction. However, the blue region of the light spectrum contains important information on the optical properties of plants, including the absorbance maxima of chlorophyll a, chlorophyll b, and carotenoids (Lichenthaler, 1987; Horton *et al*. 1996). Further advancements in the ability of hyperspectral imaging to accurately record reflectance in the blue region may potentially improve prediction accuracies for grain yield.

When predicting line performance within a site-year of interest, most site-years recorded at least one individual hyperspectral time-point with prediction accuracy equivalent or superior to prediction with markers or pedigrees, suggesting that hyperspectral reflectance phenotypes may be a useful alternative for generating predictions when markers and pedigree are not available. We also found that combining multiple time-points into hyperspectral BLUEs for a given growth stage generally did not improve accuracy beyond the average accuracy across the individual time-points. In practice, however, GY data would not be available to inform the identification of the most predictive time-points. Therefore, phenotyping at multiple time-points throughout the growing season to develop multi-time-point BLUEs may be an effective approach to buffer against time-points with low prediction accuracy.

It is also possible that some information pertaining to plant growth and development is lost when integrating multiple time-points using means-based approaches. An alternative that may more fully capture temporal variations could be to model a growth curve based on reflectance data for each line in each site-year, as in Verhulst *et al*. (2011). A greater number of time-points than were measured in this study may be required to accurately develop growth curves. In addition, statistical methods that can integrate multiple curves from a range of hyperspectral bands may be needed. Further research in this area should be performed to compare prediction accuracies of hyperspectral reflectance-based growth curves versus those achieved in this study.

Correcting results for DTHD reduced accuracies when hyperspectral reflectance was used, but not for markers or pedigrees. These results suggest that the hyperspectral reflectance measurements are also capturing physiological parameters associated with relative maturity. The greatest reductions in accuracy were observed in the Drought and Severe Drought managed treatments, where GY is typically associated with earliness. This is consistent with Rutkoski *et al.* (2014), which found that correcting for DTHD in wheat GY reduced the prediction accuracies of multivariate GS models integrating NDVI and canopy temperature measurements with markers and pedigrees. The CIMMYT bread wheat breeding program maintains a high level of diversity for DTHD in germoplasm development due to the wide target of geographic regions.. For breeding programs with high levels of variation for DTHD, it may be advisable to perform a correction for DTHD when predicting GY using hyperspectral reflectance so as to avoid indirect selection on relative maturity.

To test our proposed prediction approaches in a multi-environment context, we developed two prediction schemes. In the first, predictions were performed within a breeding cycle for genotypes that have been evaluated under some treatments but not others. In the second, predictions were performed across breeding cycles onto genotypes that were not previously evaluated. Overall, prediction accuracies in the second scheme were lower than for the corresponding models in the first, which is consistent with previous studies showing that predicting the performance of newly developed lines is more challenging than the prediction of lines that have been evaluated in correlated environments (Burgueño *et al.* 2012; Crossa *et al.* 2014).

If markers or pedigrees are not available, our results showed that predicting GY with hyperspectral reflectance alone in single-kernel models could provide similar results in terms of accuracy. It should be noted that this study was conducted at the yield trial stage of the breeding program when families contain fewer full-sibs and the variance due to Mendelian sampling is low. As pedigrees do not account for Mendelian sampling, it is possible that prediction with hyperspectral reflectance may be more advantageous than pedigree-based GS at earlier stages of the breeding program in which families contain greater numbers of full-sibs, though further study is needed to assess the ability of hyperspectral reflectance to distinguish within-family variation.

The optimal prediction accuracies were achieved by building combined models that used markers or pedigrees to model the genetic main effects and hyperspectral reflectance to model the *G* × *E* interactions. However, in prediction across breeding cycles within a managed treatment, the improvements in accuracy with the addition of markers or pedigrees were marginal when compared to the corresponding single-kernel hyperspectral reflectance models. This result was similar to Montesinos-Lopéz *et al.* (2017b), which found that the addition of markers or pedigrees did not significantly improve accuracy. The modest increases observed when predicting across breeding cycles within a managed treatment may not justify the added investment of marker genotyping.

When considering the optimal developmental growth stages during which to collect hyperspectral reflectance data, there were some differences in accuracy among growth stages when predicting with single hyperspectral phenotyping time-points within a site-year. Overall, the time-points from the HEAD and GF stages showed slightly higher accuracies than those collected during the VEG stage. However, when predicting across site-years, the advantage of phenotyping later in the season was reduced, particularly for the multi-kernel models that combined markers or pedigrees with hyperspectral reflectance, where most often there was no significant difference in accuracy among the different growth stages. Montesinos-Lopéz *et al.* (2017a) observed clear optimal time-points for hyperspectral phenotyping in a multi-environment context.

However, their study represented a balanced scenario in which all lines in all environments were phenotyped an equal number of times, and individual phenotyping time-points could be used to predict across environments. While classifying time-points according to the predominant growth stage at the time of phenotyping represents a simple and efficient method for predicting across site-years, it is somewhat difficult to compare results between growth stages. The numbers of time-points observed for each growth stage were not consistent within and across site-years. In addition, the growth stage classifications used were dependent on the predominant growth stage of the site-year and did not reflect the variation in phenology within the site-year at the time of phenotyping. These challenges may warrant further investigation into multi-environment prediction when HTP datasets are unbalanced in the number of time-points observed and at which stage of crop development those observations were recorded.

While our results suggest that hyperspectral reflectance data have the potential to add value to a breeding program by providing accurate GY predictions, the data used in this study were collected on large plot sizes that are suitable for measuring GY *per se*. While our approach may provide breeding programs with GY predictions earlier in the growing season, enabling more efficient allocation of resources at harvest, a potentially greater benefit could come from utilizing this method for smaller plot sizes where there is limited seed available to replicate and reliably assess yield. To address this, we are currently evaluating the use of aerial HTP at the early generation stage when measuring GY is not feasible.

The integration of highly dimensional data, such as hyperspectral reflectance, into relationship matrices for use in GBLUP has several advantages. First, there is no additional development in statistical software required to implement the proposed models. Many options for fitting GBLUP such as “rrBLUP”, “GAPIT”, and “BGLR” are currently available and are being increasingly used. The models proposed here can be readily implemented in these or other existing software without requiring additional user training. In addition, with the development of improved hyperspectral sensors that register reflectance at wavelengths ranging from 400 to 2500 nm, it is likely that the number of data points available for prediction will continue to increase. By using the relationship matrix-based approach to GBLUP that we have proposed, these increases in data dimensionality will neither influence the complexity of the GBLUP model, nor will they increase the computation time of the model itself. This approach could also be useful for integrating different types of highly dimensional phenotypes for prediction, such as ionomics and metabolomics data (Riedelsheimer *et al.* 2012). Rincent *et al.* (2018) recently showed that relationship matrices derived from near infrared spectroscopy absorbance of winter wheat grains and powdered leaf tissue between 400 and 2500 nm provided yield prediction accuracies that were superior to marker-based GBLUP approaches. Further research should be performed to assess the potential for this relationship matrix-based approach to be applied to other forms of biological data.

## CONCLUSION

In this study, we have proposed a multi-kernel GBLUP model that uses genomic marker-, pedigree-, and hyperspectral reflectance-derived relationships matrices to model the genetic main effects and G × E interactions across environments within a bread wheat breeding program. We have shown that deriving relationship matrices from hyperspectral reflectance phenotypes can effectively predict GY in wheat within and across managed treatments and breeding cycles. Accuracies when testing single-kernel models using hyperspectral reflectance data alone are similar to those achieved with markers or pedigrees. Our results also show that combining markers/pedigrees with hyperspectral reflectance data in multi-kernel models can increase accuracies over single-kernel approaches, but in some prediction scenarios, these increases were modest. We also suggested a method for addressing the issue of unbalanced HTP datasets involving the classification of time-points according to the predominant developmental growth stage observed at the time of phenotyping. Accuracies of multi-kernel models were roughly equivalent irrespective of the growth stage in which hyperspectral phenotyping was performed. Further research on how best to leverage multi-temporal phenotypes when the amount of data differs across site-years is needed. The methods we have proposed provide a simple and computationally efficient approach for integrating highly dimensional aerial HTP information into genomic selection and should be tested on other forms of high dimensional biological data.

## Supporting information

## ACKNOWLEDGEMENTS

We are thankful to the National Science Graduate Research Fellowship under (Grant No. DGE-1650441) and the U.S. Borlaug Fellowship in Global Food Security for supporting the graduate studies of Margaret Krause. The work presented here was supported in part by the Delivering Genetic Gain in Wheat project, supported by aid from the U.K. Government’s Department of International Development (DFID) and the Bill & Melinda Gates Foundation (OPP113319). Partial support was also provided by the Agriculture and Food Research Initiative Competitive Grants 2011-68002-30029 (Triticeae-CAP) and 2017-67007-25939 (Wheat-CAP) from the USDA National Institute of Food and Agriculture. Finally, this work was also made possible through support provided by Feed the Future through the U.S. Agency for International Development, under the terms of Contract No. AID-OAA-A-13-0005. The opinions expressed herein are those of the author(s) and do not necessarily reflect the views of the U.S. Agency for International Development.

