## Supplementary material for "Hyperspectral Reflectance-Derived Relationship Matrices for Genomic Prediction of Grain Yield in Wheat"

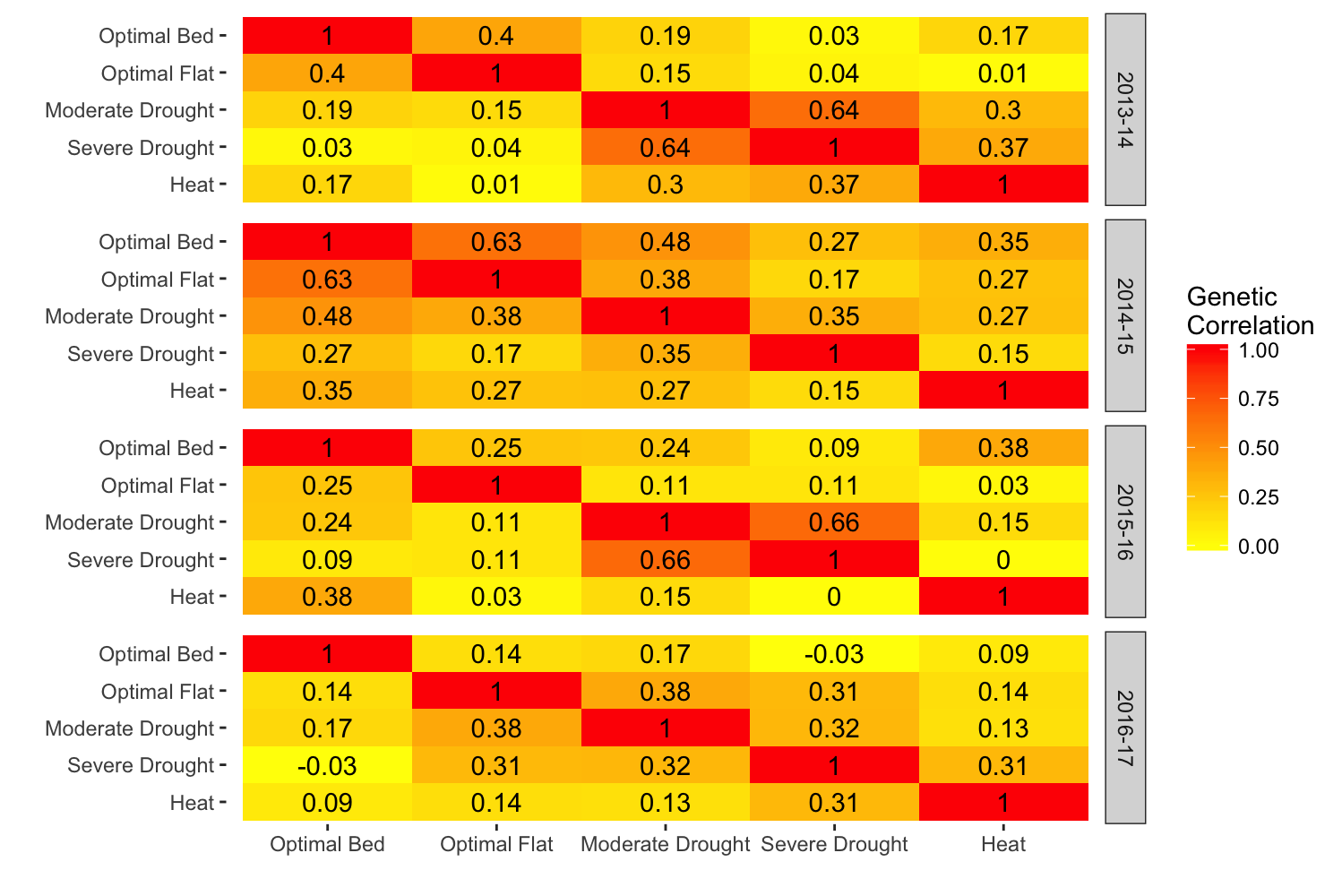

**FIGURE S1** Correlations for grain yield between managed treatments within breeding cycles

For each breeding cycle, the correlations for grain yield between managed treatments are reported. Strong and weak correlations are shown in shades of red and yellow, respectively. Intermediate correlations are shaded in orange.
