## Supplementary material for "Hyperspectral Reflectance-Derived Relationship Matrices for Genomic Prediction of Grain Yield in Wheat"

**TABLE S1** Prediction strategies used for assessing model accuracy

**A. WITHIN SITE-YEAR**

|  | **2013-14** | | **2014-15** | **2015-16** | **2016-17** |
| --- | --- | --- | --- | --- | --- |
| **Optimal Bed** | TRN | TST |  |  |  |
| **Optimal Flat** |  | |  |  |  |
| **Moderate Drought** |  | |  |  |  |
| **Severe Drought** |  | |  |  |  |
| **Heat** |  | |  |  |  |

**B. WITHIN BREEDING CYCLE/ACROSS MANAGED TREATMENTS**

|  | **2013-14** | | **2014-15** | **2015-16** | **2016-17** |
| --- | --- | --- | --- | --- | --- |
| **Optimal Bed** | TST | TRN |  |  |  |
| **Optimal Flat** | TRN | |  |  |  |
| **Moderate Drought** | TRN | |  |  |  |
| **Severe Drought** | TRN | |  |  |  |
| **Heat** | TRN | |  |  |  |

**C. ACROSS BREEDING CYCLES/WITHIN MANAGED TREATMENT**

|  | **2013-14** | | **2014-15** | **2015-16** | **2016-17** |
| --- | --- | --- | --- | --- | --- |
| **Optimal Bed** | TST | TRN | TRN | TRN | TRN |
| **Optimal Flat** |  | |  |  |  |
| **Moderate Drought** |  | |  |  |  |
| **Severe Drought** |  | |  |  |  |
| **Heat** |  | |  |  |  |

These tables represent an example of how phenotypic data records were partitioned and assigned to training and test sets for the three prediction strategies tested. Each cell represents one of the 20 site-years tested. Records allocated to the training set are indicated by TRN, while records assigned to the test set are indicated by TST. Within site-year prediction was performed considering records within a single site-year only. In this example, 80% of the records from the 2013-14 site-year were assigned to the TRN set, while the remaining 20% were assigned to the TST set. Within breeding cycle/across managed treatment prediction was performed within a single breeding cycle but across the five managed treatments. In this example, the Optimal Flat, Moderate Drought, Severe Drought, and Heat treatments from the 2013-14 breeding cycle were assigned to the TRT set, in addition to 20% of the records from the 2013-14 Optimal Bed site-year. The remaining 80% of records from the 2013-14 Optimal Bed site-year were assigned to the TST set. Across breeding cycles/within managed treatment prediction was performed across the four breeding cycles but within a single managed treatment. In this example, the Optimal Bed treatments from the 2014-15, 2015-16, and 2016-17 breeding cycles were assigned to the TRN set, in addition to 20% of the records from the 2013-14 Optimal Bed site-year. The remaining 80% of records from the 2013-14 Optimal Bed site-year were assigned to the TST set. For prediction of each site-year with each prediction strategy, 20% TRN-TST partitions were implemented.
