## Supplementary material for "Hyperspectral Reflectance-Derived Relationship Matrices for Genomic Prediction of Grain Yield in Wheat"

**TABLE S2** Summary statistics of grain yield data for each of the 20 site-years tested

| **BREEDING CYCLE** | **MANAGED TREATMENT** | **MEAN**  **(t ha^-1^)** | **STANDARD DEVIATION (t ha^-1^)** | **RANGE**  **(t ha^-1^)** | **BROAD SENSE HERITABILITY** |
| --- | --- | --- | --- | --- | --- |
| 2013-14 | Optimal Bed | 6.08 | 0.51 | 3.38-7.62 | 0.71 |
|  | Optimal Flat | 6.03 | 0.61 | 3.82-7.60 | 0.58 |
|  | Moderate Drought | 3.68 | 0.30 | 2.33-4.64 | 0.85 |
|  | Severe Drought | 2.28 | 0.43 | 0.29-3.31 | 0.94 |
|  | Heat | 2.08 | 0.46 | 0.05-3.00 | 0.93 |
| 2014-15 | Optimal Bed | 5.61 | 0.48 | 3.56-6.78 | 0.82 |
|  | Optimal Flat | 5.72 | 0.52 | 3.74-6.97 | 0.74 |
|  | Moderate Drought | 4.52 | 0.36 | 2.96-5.69 | 0.79 |
|  | Severe Drought | 3.80 | 0.51 | 1.05-5.16 | 0.67 |
|  | Heat | 2.73 | 0.35 | 1.51-4.38 | 0.84 |
| 2015-16 | Optimal Bed | 7.15 | 0.37 | 5.89-8.29 | 0.75 |
|  | Optimal Flat | 6.97 | 0.48 | 4.68-8.74 | 0.68 |
|  | Moderate Drought | 3.27 | 0.33 | 0.67-4.04 | 0.82 |
|  | Severe Drought | 3.66 | 0.40 | 1.88-4.99 | 0.93 |
|  | Heat | 1.63 | 0.46 | 0.04-2.72 | 0.79 |
| 2016-17 | Optimal Bed | 6.51 | 0.55 | 4.26-7.80 | 0.71 |
|  | Optimal Flat | 6.28 | 0.68 | 3.09-7.96 | 0.80 |
|  | Moderate Drought | 4.84 | 0.42 | 3.30-5.97 | 0.82 |
|  | Severe Drought | 2.63 | 0.50 | 0.50-3.94 | 0.83 |
|  | Heat | 3.95 | 0.44 | 1.53-5.11 | 0.88 |
