## Supplementary material for "Hyperspectral Reflectance-Derived Relationship Matrices for Genomic Prediction of Grain Yield in Wheat"

File S1 Within site-year prediction accuracies and standard errors; File S2 Within breeding cycle/across managed treatments prediction accuracies and standard errors; File S3 Within managed treatment/across breeding cycles prediction accuracies and standard errors; TABLE S1 Prediction strategies used for assessing model accuracy; TABLE S2 Summary statistics of grain yield data for each of the 20 site-years tested; FIGURE S1 Genetic correlations for grain yield between managed treatments within breeding cycles
